# Thriving in the heat – Lysine acetylation stabilizes the quaternary structure of a Mega-Dalton hyperthermoactive PEP-synthase

**DOI:** 10.1101/2022.08.11.503304

**Authors:** Pascal Albanese, Wenfei Song, Siri van Keulen, Jeroen Koendjbiharie, Fujiet Koh, Barbara Steigenberger, Tomoko Vincent, Albert Konijnenberg, Albert J.R. Heck, Servé W.M. Kengen, Alexandre M.J.J. Bonvin, Friedrich Förster, Richard A. Scheltema

## Abstract

Over time structural adaptations enabled proteins and enzymes to have sufficient stability and flexibility to perform the basic functions of life under various environmental conditions. The catalytic cores of key metabolic enzymes of hyperthermophilic archaea work at a temperature range of 80-120 °C, similar to the conditions wher the earliest life forms may have thrived. Here we characterize a key enzyme of the central carbon metabolism of Pyrococcus furious, through an integrative approach combining structural mass spectrometry, cryo-electron microscopy, mass photometry and molecular modelling with molecular dynamics simulations. From our investigation, we unveil the structural organization of phosphoenolpyruvate synthase (PPSA). Its 24-meric assembly - weighing over 2 MDa - harbors flexible distal domains, whose proper functioning and coordination depends on widespread chemical acetylation of lysine residues. This non-enzymatic post-translational modification, along with other types of lysine modifications, also occurs on most other major protein complexes of P. furiosus. These modifications likely originated in the chemically favorable primordial conditions and gradually became highly specialized and enzyme-driven in more distantly related mesophiles and Eukaryotes.

---

Life has conquered nearly every habitat on the planet and, although the thermophilic origin for early organisms is still debated(Weiss et al., 2016), bacterial and archaeal species have even been isolated from marine hot environments that may retain key characteristics of the earliest life forms(Martin et al., 2008). Such habitats forced these organisms to develop extensive strategies geared towards stabilizing their cellular components, primarily proteins, to such a degree that the metabolic reactions essential to support life could be performed(Rothschild and Mancinelli, 2001). On the other hand, catalytic domains inside many proteins also require a degree of flexibility to enable their activity, requiring organisms to perform an intricate balancing act between stability and flexibility. Hyperthermophilic Archaea, organisms with an optimal growth temperature exceeding 80 °C, perform this balancing act on a daily basis and hence have been of major interest for biotechnology, and many enzymes have been isolated, characterized and exploited for decades owing to their exquisite stability(Vieille and Zeikus, 2001). There however is no clear single factor determining the increased (thermal) stability of proteins. This is highlighted by numerous studies uncovering the importance of noncovalent stabilization strategies like hydrophobic(Lieph et al., 2006) and ionic interactions(Karshikoff and Ladenstein, 2001), deletion of exposed loop regions(Thompson and Eisenberg, 1999), metal binding(Kardinahl et al., 2000), and covalent disulfide bonds(Jorda and Yeates, 2011). Besides stability, most proteins engage in protein-protein interactions to enable functionalities that exceed the sum of the individual parts, and it is becoming increasingly clear that this functional interactome is dynamically wired and rewired through alterations of protein post-translational modifications (PTMs)(Bludau and Aebersold, 2020). These molecular switches controlling protein function are well known to play pivotal roles in mesophilic organisms and phosphorylation(Cohen, 2000) & acetylation(Choudhary et al., 2009) are for example heavily investigated. Their role in Archaea and in particular their significance in thermophilic environments are however still largely unexplored. Lysine acetylation occurs across all domains of life and can spontaneously form or enzymatically deposited. These modifications fulfill diverse roles beside the established and conserved transcription regulation via *e*.*g*. histone modifications(Yang, 2004). Non-enzymatic acetyl coenzyme A (acetyl-CoA)-dependent acetylation was shown to play a role in mitigating metabolic and environmental stress in prokaryotes(Hentchel and Escalante-Semerena, 2015). This is achieved through direct regulation of enzymes of the central carbon metabolism(Wang et al., 2010), which also occurs through non-enzymatic coupling(Nakayasu et al., 2017). Recent proteomics studies found that up to 20-25% of the total proteome in Archaea contain acetylated lysine residues, highlighting a role for this PTM in carbon metabolism(Cao et al., 2019; Liu et al., 2017). The question remains whether this is the product of specific regulation, or a byproduct of spontaneous chemical reactions that occur when environmental conditions and intermediate metabolic substrates are highly abundant as occurs for example in metabolic enzymes of mitochondria(Baeza et al., 2016).

All biomass production on Earth relies on a combination of a handful of precursor metabolites driving carbon metabolism. Of this limited set, phosphoenolpyruvate (PEP), pyruvate and oxaloacetate are at the intersection of glycolysis and the tricarboxylic acid cycle forming the so-called PPO-node(Koendjbiharie et al., 2020). Variations within this node are pivotal to metabolic plasticity, ultimately reflecting metabolic adaptation to extreme ecological niches. One key enzyme in the PPO-node is PEP synthase (PPSA, EC 2.7.9.2), which catalyzes the following reaction(Flamholz et al., 2012):

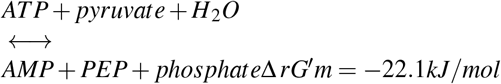

The reaction consists of two intermediate steps. (1) A phosphate group is transferred from ATP in the Nucleotide Binding Domain (NBD) to an intermediate phosphorylated amino acid of the mobile Central Domain (CD). (2) Then – similarly to the functionally and structurally related enzyme pyruvate phosphate dikinase (PPDK, EC 2.7.9.1) (Lim et al., 2007) – a large displacement of the CD domain carries the phosphate to the final acceptor pyruvate in the PEP/Pyruvate Binding Domain (PBD)(Minges et al., 2017). Although the reaction thermodynamically favors the gluconeogenic direction toward PEP, some PPSA complexes have been shown to play a central role in glycolytic conversion of PEP to pyruvate, with possible implications in the regulation of ATP/GTP/NADPH ratios(Imanaka et al., 2006). Despite its central role in many prokaryotes, structural information on this molecular machine is largely lacking. To date, a single dimeric PPSA X-ray crystal structure is available from the pathogenic bacterium *Neisseria meningitidis* (PDB: 2ols), and some examples are available for PPDK enzymes from bacteria(Herzberg et al., 1996) and eukaryotes(Cosenza et al., 2002). From hyperthermophilic archaea, two PPSA’s have so far been described using different techniques. A large 24-meric homo-complex from Staphylothermus marinus was described by Harauz and co-workers in the 90’s – pioneering applications of single particle electron microscopy(Cicicopol et al., 1994, 1999; Harauz et al., 1996; Li et al., 2000) – which could however only roughly describe the arrangement of the core of the complex due to the flexibility of the outer CD and NBD domains. Additionally, biochemical and functional assays were reported for PPSA from Pyrococcus furiosus describing 16-meric and octameric complexes, of which only the latter was found to be catalytically active(Hutchins et al., 2001). These experiments were however performed in the presence of reducing agents to allow anaerobic conditions during the first purification steps, thus with possible effects on complex integrity.

In this work, we purify and characterize a catalytically active 24-meric PPSA homocomplex from *Pyrococcus furiosus*. A structural model of the complete enzyme was obtained following an integrative structural biology approach: high resolution mass spectrometry pinpointed the PTMs; cryo-electron microscopy (EM) revealed the structure of the stable C-terminal core complex and the variable positioning of the mobile CD domain; crosslinking mass-spectrometry (XL-MS), co-evolutionary analyses and structural modelling were combined with molecular dynamics simulations (MD) to determine the role of PTMs on the arrangement of the flexible NBD domain and its sub-stoichiometric interactors. The striking structural features stabilizing the PPSA complex are novel findings that have implications for rational protein design and also shed light on the origin of widespread spontaneous acetylation in thermophilic organisms.

## Results

### Widespread lysine acetylation in high molecular-weight protein complexes

*P. furiosus* cells of two independently grown cell batches (strain DSM 3638, see Methods for details) were lysed in mild conditions and fractionated by sequential centrifugation to isolate the soluble proteome (Supplementary Figure 1). Of the theoretical 2036 coding genes(Bridger et al., 2012) 1228 proteins could be identified in a single-shot experiment (Supplementary Table 1), a number comparable with previous studies(Wong et al., 2013). To extract high-molecular weight (MW) complexes, the soluble proteome was further fractionated by either sucrose gradient ultracentrifugation (SG) or size-exclusion chromatography (SEC) (Supplementary Figure 2). By supplementing these enriched fractions, we were able to increase the number of identified proteins to 1,435 (Supplementary Table 1). Intensity-based absolute quantification (iBAQ) values(Cox and Mann, 2008) and SDS-PAGE gels obtained from the different SG and SEC fractions clearly showed that many intact multi-protein complexes were purified (Figure 1A, Supplementary Figure 2). The set of protein complexes comprised many known actors, among others the Heat Shock Protein (Hsp) 60 thermosome, the 50S and 30S ribosomes, and the CRISPR/Cmr complex. A surprisingly abundant protein complex is the PPSA enzyme, which far exceeds the abundance of up- and down-stream metabolic enzymes of the PPO-node. Namely, the phosphoenolpyruvate carboxylase (Q8TZL5) and carboxykinase (Q8U410) are between 300 and 1,000 times less abundant than PPSA (Supplementary Table 1). In previous studies lysine acetylation was found as a widespread modification in prokaryotes and we therefore included this modification as a variable modification in our searches, along with other previously reported lysine acylations – modifications that are most likely derived from CoA derivatives(Baeza et al., 2016). We here report a first-of-its kind analysis in which each of these modifications was identified and quantified on a proteome-wide scale simultaneously. We report a widespread occurrence of acylated lysines with 400 or more sites detected for each of them (Figure 1B), with however a prevalence of acetylation, with 1,433 total sites (Supplementary Table 1). Lysine acetylation (acK) is, however, not standing out compared to other acylations in term of total intensity in the fractionated lysates (Figure 1B,C), but it is significantly enriched in the high-molecular weight (MW) fractions, namely with MW above 400-500 kDa (Figure 1C). Our depth allowed us to determine that among the over one thousand detected acK sites, almost half were distributed among seven main protein complexes (Figure 1B). Our investigation of the possible role of these widespread lysine modifications, and specifically the acetylations, further targeted an unusually abundant metabolic enzyme, a supposedly homomeric form of PPSA (Supplementary Figure 2) identified by MS on the excised gel band with a sequence coverage of approximately 98% (Supplementary Table 1).

**Figure 1.**
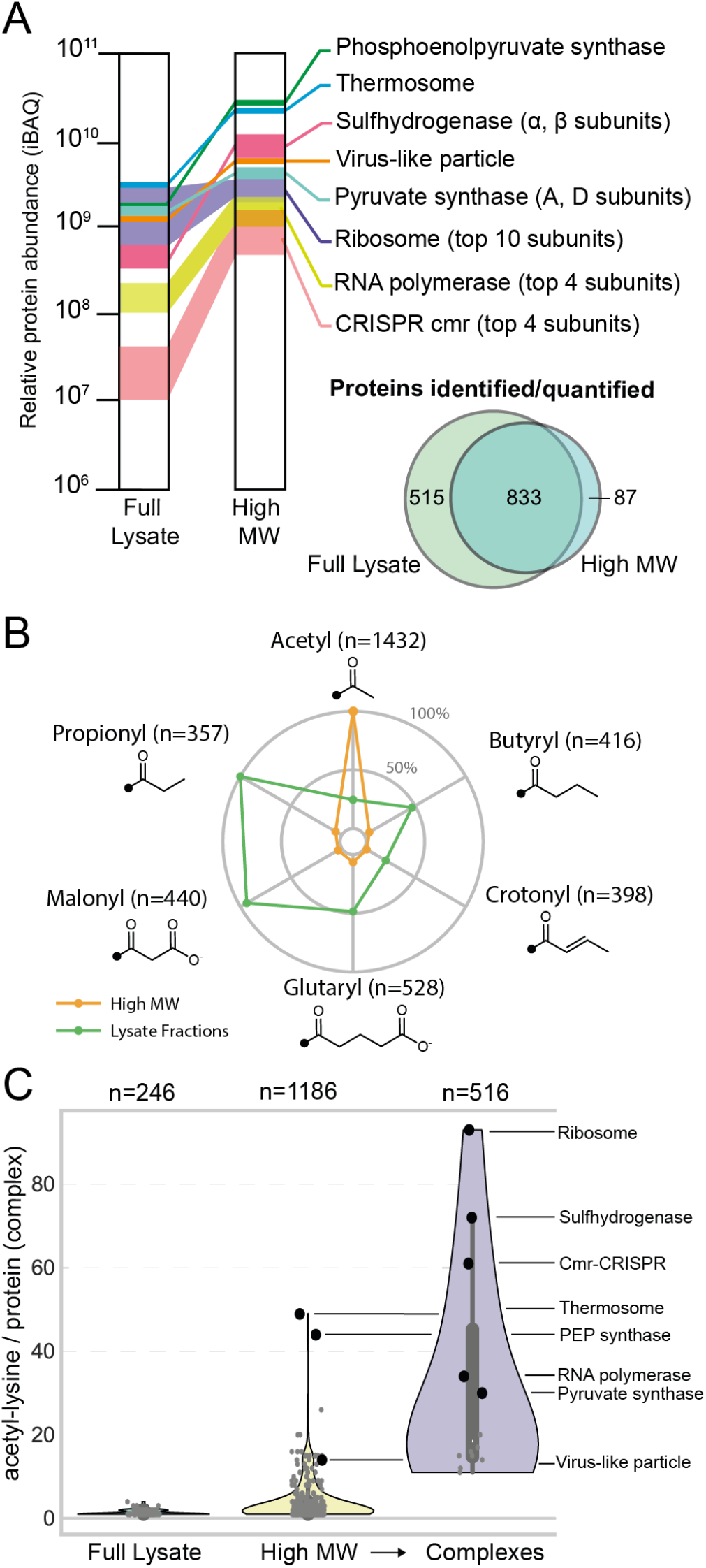
The *P. furiosus* proteome undergoes widespread acylation on Lysine residues, among which acetylation preferentially occurs in high-molecular weight complexes. Relative abundance in the proteome of the main soluble multiprotein complexes in the full lysate or enriched by SG and SEC fractionation (high MW) of the same cell lysate **(A)**. Relative abundance and distribution of the 3571 acylated lysine (acK) sites detected by bottom-up proteomics analysis on the fractionated proteome and high MW fractions **(B)**. Distribution of acK residues per protein or grouped into known protein complexes **(C)**.

### *P. furiosus* PEP synthase is a heterogeneous mega-Dalton complex stabilized by Methionine-Iron coordination

*P. furiosus* PPSA was purified by sucrose gradient (SG) ultracentrifugation as a heterogeneous mixture of high MW assemblies with a total mass ranging from 1.9 and 2.3 MDa (*>*83% of the total particles of the SG fraction), with some assemblies as low as 1.5 MDa or as high as almost 3 MDa, as determined by single particle counting in solution via Mass Photometry (MP) (Supplementary Fig 3A, Supplementary Data 2). This analysis revealed the heterogeneity of this complex in terms of mass, suggesting that different oligomerization states and/or heteromeric complexes may be present. Isolated complexes were vitrified for single-particle cryo-EM analysis, resulting in a reconstruction with an overall resolution of 2.9 Å of a 24-meric assembly, likely representing the preferred configuration of *P. furiosus* PPSA (Figure 2, Supplementary Fig 4). Residues 506 to 807 were directly modelled from the cryo-EM density, and the last C-terminal residues (808-814) were included from the fitted AlphaFold2 predicted model (see Methods for details). The cryo-EM model comprises 24 C-terminal domains assembled in 6 tetrameric modules that tethered by the planar coordination of an iron ion (detected by ICP-MS, Supplementary Table 2) by four Methionine (MET) 799 residues facing inward (Figure 2 – middle left) and further stabilized by inter-chain salt bridges (Figure 2 – bottom left). MET-Fe(II) coordination is, to our knowledge, rare in structures, but closely resembles that observed in an “ion channel” of the *P. furiosus* ferritin (PDB: 2×17, Supplementary Figure 5), suggesting that this stabilizing strategy is shared by other *P. furiosus* protein complexes and, possibly, by other organisms living in anoxic iron-rich environments. Every PPSA unit also has a dimeric interface with a PPSA counterpart of a neighboring tetrameric module, which is strengthened by hydrogen bonds (Figure 2 – middle right) and a strongly conserved arrangement that resembles the one from bacterial and eukaryotic (plant) PPDK enzymes (Supplementary Figure 6). The PBD folding of PPSA/PPDK is also remarkably conserved among the 3 kingdoms of life, with a root mean square deviation of the backbone below 4.2 Å of the pruned atoms and the same arrangement of the residues facing the PBD catalytic pocket (Supplementary Figure 5).

**Figure 2.**
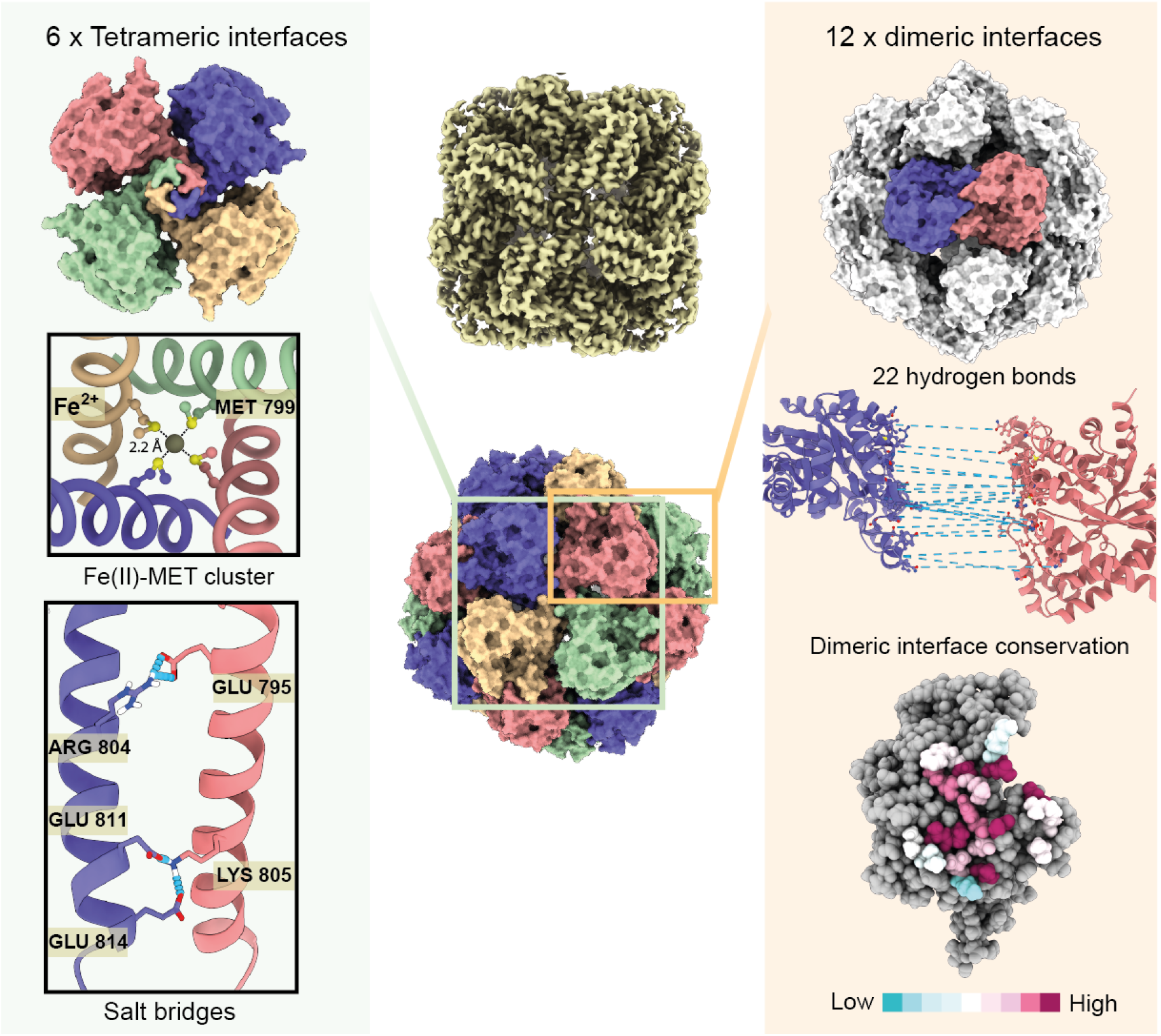
Structural arrangement of the PEP-synthase stable core. Cryo-EM density map and surface representation of the 24 subunits organized into 6 tetrameric interfaces stabilized by metal coordination and salt bridges (left panel). Each subunit also has a conserved dimer interface in a neighboring tetramer (right panel), contact residues (¡2Å) between monomers are color coded based on conservation scores (Ashkenazy et al., 2016).

### PPSA 24-mer oligomerization is shared by other archaea and needed for thermal stability

The functional unit for PPSA is a dimer based on homology with PPDK (Ciupka and Gohlke, 2017). These were however mostly studied in plants and few bacteria. The activity of both the gluconeogenic and glycolytic reactions were measured (Supplementary Table 3 and Supplementary Figure 3) showing rates comparable to similar enzymes. Under reducing conditions (10mM TCEP), the 24-mer oligomerization is disrupted in favor of a tetrameric assembly, but only half of the catalytic efficiency is retained at 80-90 °C (Supplementary Figure 3). At higher - but still lower than physiologically relevant - protein concentrations, this residual activity under reducing conditions disappears as proteins aggregate (depicted by the decreasing number of total particles counted at 80-90 °C) (Supplementary Fig 3C). The conditions at which this occurs is strongly indicative of temperature- and/or concentration dependent aggregation, as this drop does not occur at lower temperatures (25-60 °C). Interestingly, we did not observe the dimeric state of PPSA in any of the conditions tested, suggesting that the tetrameric assembly is the minimal catalytic unit of *P. furiosus* PPSA. At catalytically relevant temperatures and concentrations, however, the tetramers aggregate (Supplementary Figure 3) indicating that their assembly into a full 24-mer is required.

As the structural feature allowing their stabilization revolves around MET-Fe(II) coordination, we combined the motif distribution in organisms containing a methionine in this position and flanking charged amino acids with a phylogenetic comparison of PPSA homologous sequences in bacterial and archaeal taxa (Supplementary Data 4). The 2.25 MDa PPSA assembly previously characterized in *S. marinus* (Crenarchaeota) shares the same oligomerization state (Harauz et al., 1996; Li et al., 2000) and deletion of the 20 C-terminal residues containing the conserved MET-799, prevented its oligomerization (Cicicopol et al., 1999). Crenarchaeota (*S. marinus*) and Euryarchaea (*P. furiosus*) seem to diverge very early based on the PPSA sequence, sharing the origin of the unrooted tree with other bacterial taxa (Supplementary Figure 7) where this motif, and possibly this arrangement, is also conserved. A clear correlation between (hyper-)thermophilic niches and the 24-meric oligomerization appears (Supplementary Figure 7). The PPSA gene was shown to be horizontally transferred from bacteria to archaea early in their evolutionary history (Nelson-Sathi et al., 2015) and may have subsequently evolved in the functional PPDK-like dimeric arrangement at milder temperatures.

### The cryo-EM data predominantly highlights a enzymatic resting state of PPSA

PPDK enzymes require large conformational changes to shuttle the phosphate group back and forth between ATP in the NBD and PEP in the PBD, which in *P. furiosus* PPSA is located within the stable core solved by cryo-EM (Figure 3). This transfer is enabled in mesophilic PPDK enzymes through the CD domain that undergoes a “swiveling” mechanism (Lim et al., 2007). As the isolated PPSA is catalytically active around the expected mass, a similar mechanism involving the CD domain – with a large degree of flexibility – can be anticipated and indeed several low resolution densities denoting multiple conformations were found in the Cryo-EM 3D classes (Figure 3A and Supplemental Figure 4). The position of the CD domain favored by the EM data was determined, at a resolution of 7.1 Å, to be located on top of the PBD domain and at a distance from the PBD catalytic sites too distal to enable direct transfer (Figure 3B,C and Supplementary Figure 8). The homology model for the CD domain, upon further energy minimization after fitting into the cryo-EM density, revealed that this PBD-CD interaction is stabilized by three inter-domain salt bridges (Figure 3C). This representative inactive state is hereafter called the “resting” state. As the conformational switch could however be induced at physiologically relevant temperatures – reaching full operability in both gluconeogenic and glycolytic directions (Supplementary Table 1 and Supplementary Figure 3) – we also modelled the catalytically active states.

**Figure 3.**
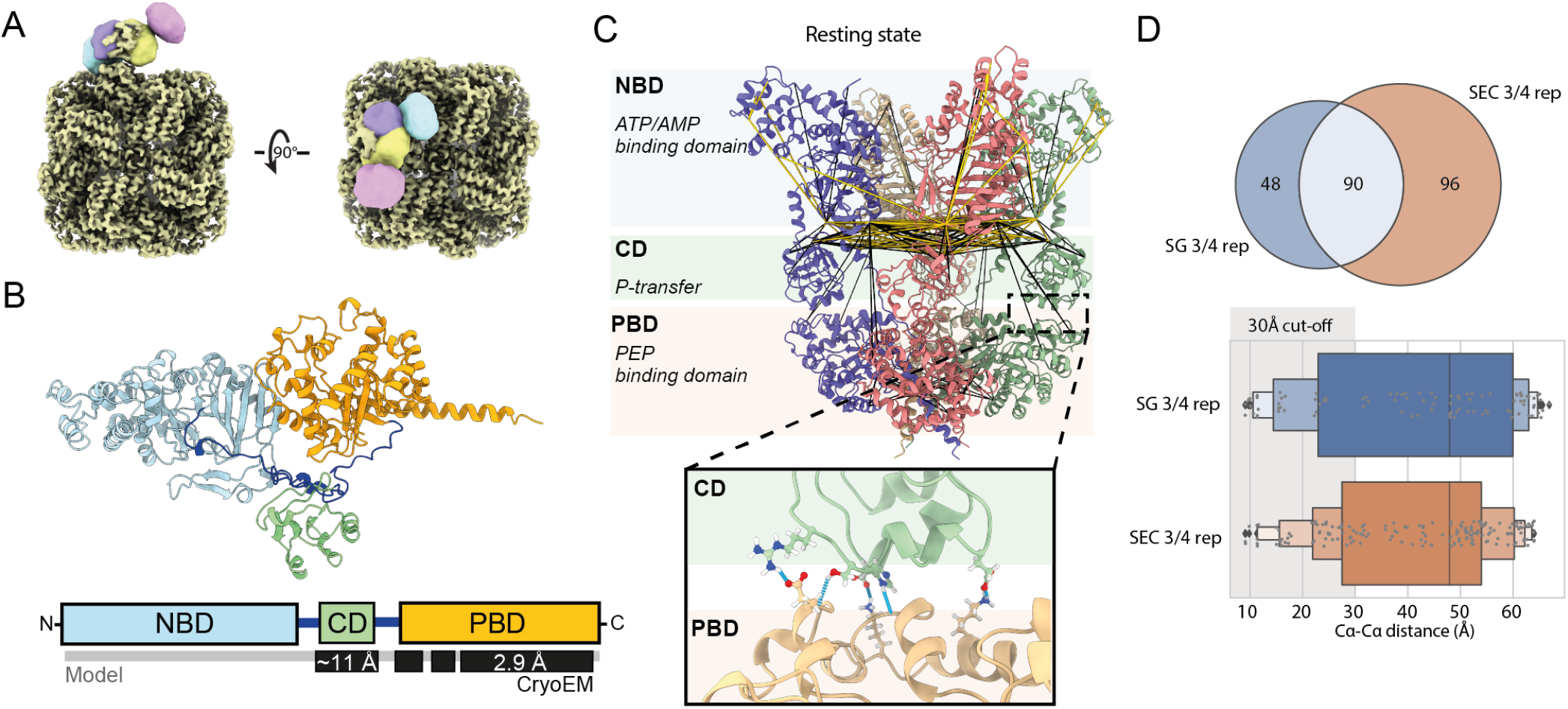
Integrative modelling and focused refinement of the mobile and N-terminal domains unveil the required flexibility for catalysis. Different possible orientations of the central domain (CD) around the PEP-binding domain (PBD) from focused refinement of the cryo-EM data **(A)**. Structural model of the PEP synthase monomer colored according to the functional domains **(B)** with experimentally derived and predicted sequence portions indicated along with their highest cryo-EM resolution. Assembly of the PPSA “rest” state, the representative conformation in our sample based on Cryo-EM, XL-MS and co-evolution sequence analysis data **(C)**. Hydrogen bonds stabilizing the CD domain in the “rest” position are shown in the inset. Euclidean distances between the backbone carbons of reproducibly crosslinked lysine residues from SG and SEC samples are shown on the PPSA “rest” state model **(D)**.

### Three distinct PPSA conformations form during the catalytic cycle

Functional measurements showed that the purified complex has full catalytic capabilities in both the gluconeogenic and glycolytic direction at 80°C (Supplementary Table 3), with efficiencies comparable to enzymes of similar class (Hutchins et al., 2001; Lim et al., 2007). To investigate the structural impact of our experimental findings, a combination of our experimental results and (structural) bioinformatics tools were employed to model three all-atom conformations of PPSA in the tetrameric form to assess the conformational range of the enzyme (from resting state to near-catalytic conformations) and to investigate the location of the 14 acetylated lysines (see Methods) within the context of the PPSA 24-mer.

The first state we modelled was the CD-NBD state for which we used HADDOCK(De Vries, Van Dijk, and Bonvin 2010), MODELLER(Eswar et al., 2006) and PEP-FOLD(Lamiable et al., 2016). Distance restraints included in the protein-protein docking calculations performed with HADDOCK were extracted from an intra-molecular co-evolution analysis(Hopf et al. 2019) with false positives removed by DisVis (van Zundert et al. 2017)(see Methods). The XL-MS and cryo-EM results were used solely to validate the resulting PPSA tetramer models. Subsequently, a model of the CD-PBD state was constructed by using a PPDK structure(Lim et al., 2007) in which CD is in close proximity to PBD (ATP/pyruvate conversion cycle) as a template (see Methods) to adapt the predicted CD-NBD state to the CD-PBD conformation using MODELLER(Eswar et al., 2006). The “resting” state was constructed by using the CD-PBD state in combination with the cryo-EM data of the CD position on top of the PBD domain to reorient the domains and to model the linker between the CD and the PBD regions. All three predicted structures (CD-NBD, CD-PBD, resting) were energy minimized and acetylated through HADDOCK and their quality was assessed by checking their overlap with the XL-MS and cryo-EM data as well as with the residue positioning required for ATP and pyruvate conversion.

From these models it is evident that the long linkers between CD-NBD and CD-PBD play a crucial role in the conversion cycle as they enable the relocation of CD to interact with NBD or PBD and initiate the ATP/pyruvate or PEP/AMP reaction mechanisms (see Supplementary Note 1 for details). Even though the presence of these flexible regions is not surprising for mesophilic organisms, it is likely a liability at the extreme conditions (100°C) in which this enzyme operates due to a potential increase in dynamics induced by the elevated temperatures. As the biochemical and structural proprieties of enzymes appeared to largely depend on PTMs that extensively occur in *P. furiosus*, we used high-resolution MS proteomics to investigate the presence and the role of phosphorylation and acetylation on PPSA structural dynamics.

### Threonine 440 is the phosphorylated intermediate

Phosphorylation of a residue within the CD domain is a required intermediate step to transfer the reducing equivalents to and from PEP between the two spatially distant catalytic sites in PBD and NBD, respectively. Within our shotgun proteomics measurements, we identified 23 fragmentation spectra that support phosphorylation in the previously reported region, even without specific enrichment for phosphorylated peptides and with an overall occupancy of *>*60% (Supplementary Table 1 and Supplementary Data 3). Due to the presence of three consecutive putative sites (Thr-440, Ser-441 and His-442), all equally conserved, it was necessary to carefully evaluate the localization probability for each position. Clearly, His-442 is the least likely based on this analysis (Figure 4A), although it cannot be excluded this is driven by the lower stability reported previously for histidine phosphorylation that is further exacerbated by the elevated temperatures(Potel et al., 2018). Considering however that the orientation of the CD in the resting state buries His-442 in the structure and Ser-441 likely is involved in hydrogen bonding, leaves Thr-440 as the preferred phosphorylation site (Figure 4A). Although it was previously reported that phosphorylation occurs on a histidine residue of the PPDK CD domain through indirect biochemical assays(Spronk et al., 1976), recent proteomics studies on PPDK extracted from plant also report homologous threonine and serine residues as the phosphate carriers(Chen et al., 2014; Reiland et al., 2009).

**Figure 4.**
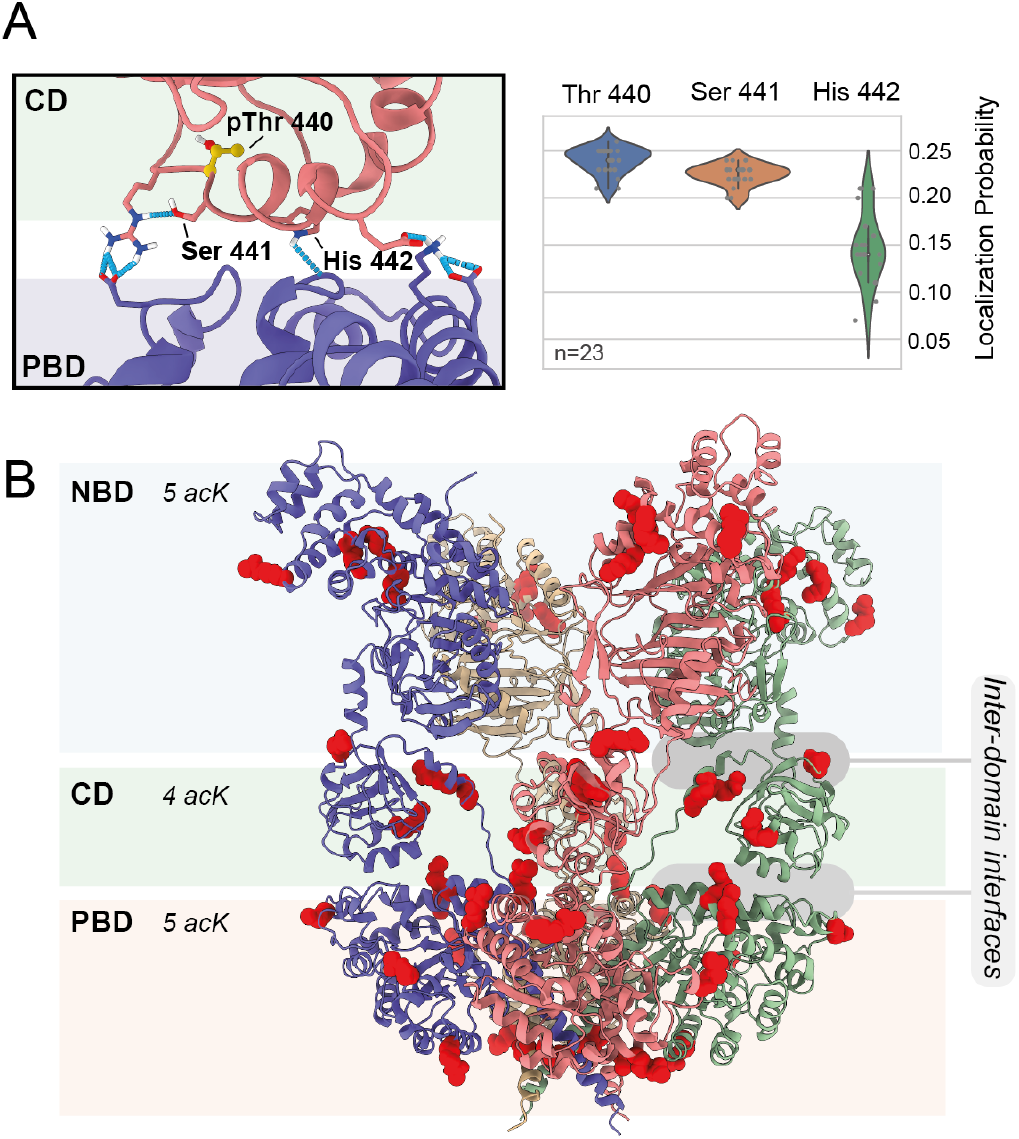
Functional post-translational modifications of PPSA. Phosphorylated residues of the CD domain are close to the interface with the PBD and localization probabilities for each site were evaluated on 23 spectra **(A)**. Acetylated lysine (acK) sites with high occupancy (Supplementary Figure 10) are highlighted in red on the “rest” state PPSA model **(B)**.

### Spontaneous lysine acetylation increases the heterogeneity of the PPSA pool

By modifying the net charge of lysine residues, acylations can have a profound effect on the overall function of the protein. In this study, we identified numerous acylated lysine sites located either on homomeric or heteromeric complexes of which acetylation is the dominant form (Figure 1B). PPSA is not deviating from this trend, with 51 out of the 59 lysines found in acetylated form 16 of which at high stoichiometry (Figure 4B and Supplementary Figure 10), but without a specific sequence motif (Supplementary Figure 14). This was expected, as extensive acetylation machinery, excluding a few specific targets for chromatin remodeling like the protein Alba, is missing(Bell et al., 2002). Thus, the existence of such a diverse acetylome can only be driven by a non-enzymatic transfer within the cellular environment. A mechanism to achieve this, known from mitochondria involves cysteine residues located within 10 Å as carriers for the acetyl group derived from acetyl-CoA (James et al., 2017; Wagner and Payne, 2013). Strikingly, cysteines were in fact reported to occur less frequently in thermophilic organisms due to their elevated and deleterious reactivity at high temperatures (Vieille and Zeikus, 2001). In PPSA, a total of five cysteines are present, of which only Cys756 is engaged in the catalysis and is located in the PBD core (Supplementary Figure 5). The remaining “accessory” Cys746 and Cys454 of the PBD and CD domains, respectively, are between 10 and 25 Å away from highly acetylated lysines suggesting a similar mechanisms of spontaneous acetylation (Supplementary Figure 10B). To strengthen this observation, we mapped the acK sites detected by MS on complexes with known 3D structure – the lysyl amino-peptidase complex (homo 12-mer, PDB: 4×8i) from *P. furiosus* and the aminopeptidase TET2 from *P. horikoshii* (PDB: 6f3k). Within these complexes, only one cysteine is present per monomer and it is located around 18-20 Å (between 10 and 25 Å) from acetylated lysines (Supplementary Figure 10), further supporting the possible role of cysteine-mediated spontaneous acetylation in *P. furiosus*. An enrichment in lysine residues on the exposed surface of hyperthermophilic proteins was already reported, but was linked to an increased amount of ion pairs at the surface – possibly increasing the stability of coulombic interactions at elevated temperatures(Cambillau and Claverie, 2000) as also suggested by increasing in vitro lysine acetylation rates of purified enzymes (Shaw et al. 2008). Acetylation also preferentially occurs at solvent accessible residues, neutralizing lysine charge, which can be in turn advantageous in the highly crowded cytosolic compartment to avoid aggregation of abundant proteins and protein complexes. An important point considering the high abundance of PPSA. Several other acylations occurring with lower stoichiometry (Supplementary Figure 9) contribute to the highly differentiated and variable pattern of PPSA chemical modifications, ultimately determining the wide range of its mass distribution. To further investigate the role of acetylation on the structural dynamics of PPSA we used coarse-grained (CG) molecular dynamics (MD) simulations.

### Lysine acetylation stabilizes the overall organization of the flexible NBD domains and allows catalysis

We performed a coarse-grained simulation of the full 24-mer PPSA model in the resting state and the CD-PBD state. During the simulation, two models – one with and one without acetylation - were run with Thr-440 phosphorylated at 90 °C in a fully hydrated state and in the presence of physiologically relevant monovalent ions (150 mM NaCl). First, we investigated two conformations of the resting state which could be possible due to interdomain linker loop flexibility. The two possible arrangements of the “resting” state are: the PBD-CD-NBD contacts interfaces are all within the same monomer (state 1), and the CD-NBD module is sitting on top of the PBD from the neighboring monomer (state 2, Supplementary Figure 11). Independently of lysine acetylation, state 2 was not a stable conformation, with the system unfolding before approximately 100ns in all 4 replicate runs (Supplementary Figure 11). We therefore extended the simulation only for state 1 (all intra-monomer interfaces) to 500ns in two replicates per acetylation state to investigate the impact of lysine acetylated within the PPSA 24-mer. The two 24-mers simulated independently for each condition (with and without acK) were used to extract the trajectories from for each of the 12 PPSA tetramers (2*6 tetramers), which were analyzed seperately (Supplementary data 5). Notably, the simulations highlight that the presence of acetylated lysine residues induce a more ordered re-arrangement of the NBD domains during MD (Supplementary data 5), that finally stabilize in most cases into a dome-shaped assembly, contrarily to the less ordered arrangement resulting from the presence of unmodified lysines (Figure 5A). This stabilization effect is evident looking at the root-mean-square deviation (RMSD) of the tetrameric NBD domains with respect to the initial “resting state” model (Figure 5B). Towards the end of the simulation, and only in the presence of acetylated lysines, we observe the sampling of functional CD-PBD contacts (Figure 5C,D), indicating that the phosphorylated CD domain migrates towards the PBD catalytic site where the enzyme performs the Pyruvate *>* PEP conversion. These findings imply that acetylation has profound implications for maintaining the NBD catalytic site in a reasonable proximity to the CD domain to promptly participate in the phosphate-transfer reaction. The outcome of the MD simulations is further strengthened by mapping on the resulting structures the distance restraints from seven independent XL-MS replicates, showing that nearly all crosslinks fall within the 30 Å cut-off (Figure 5E).

**Figure 5.**
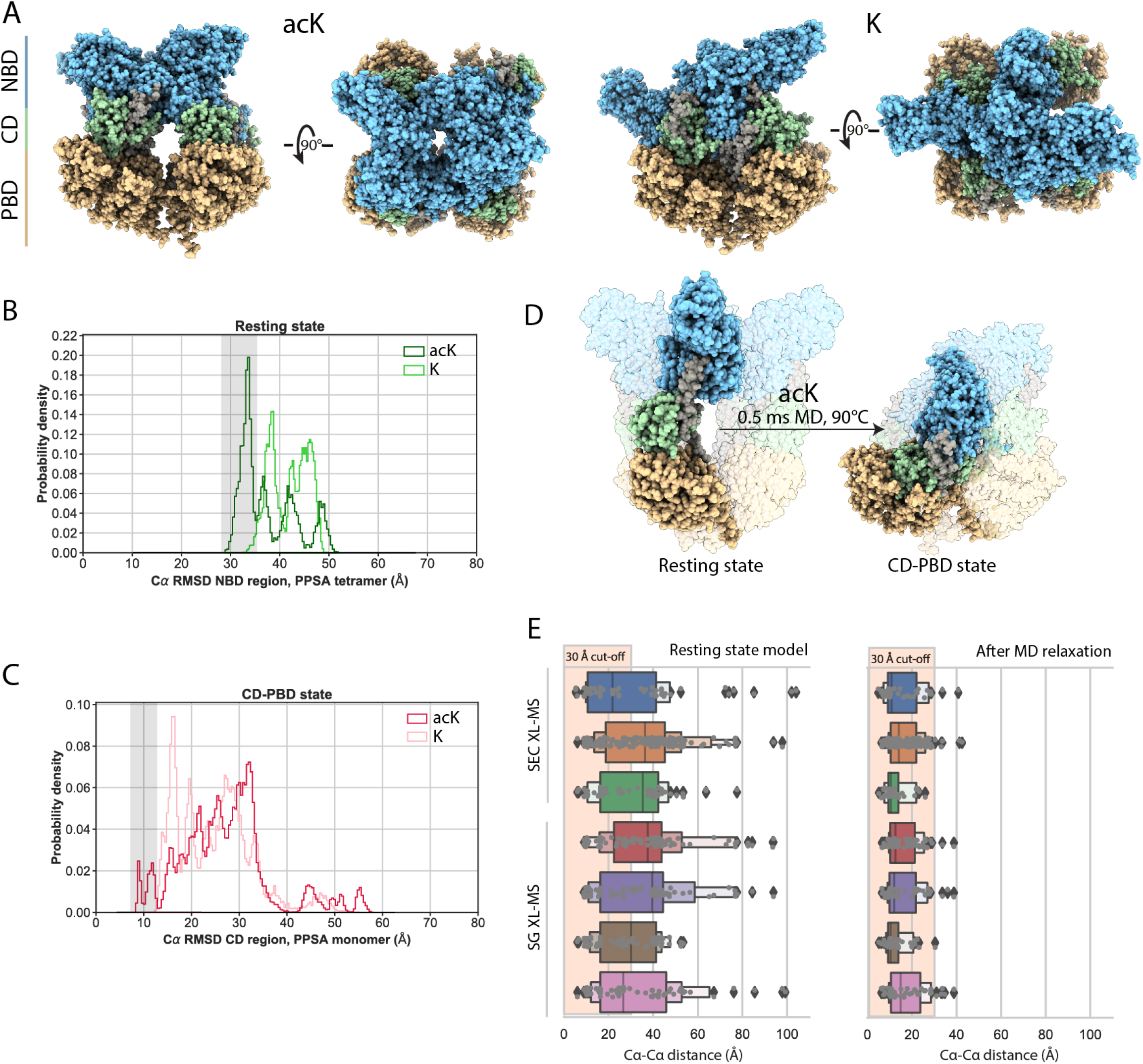
Molecular dynamics simulations unravel the stabilizing function of widespread lysine acetylation. The full 24-mer model was coarse grained with (ALY) and without (LYS) acK sites using the starting “rest” configuration, representative states of the tetramers at the end of 0.5 ms simulations are shown **(A)**. The NBD region deviates less from the starting conformation in the presence of acK, pointing to a stabilized conformation **(B)**, and the CD domain approaches the functional predicted CD-PBD state where the phosphate on THR-440 is transferred to pyruvate **(C)**. This functional transition is sampled uniquely in the presence of acK **(D)**. The shortest backbone carbons Euclidean distance before and after MD simulation between two crosslinked lysines is shown for 7 independently purified PPSA preparations obtained by SEC and SG fractionation **(E)**.

### A stably interacting Phosphoenolpyruvate carboxykinase suggests PPSA plays multiple roles in *P. furiosus* metabolism

Given the skewed mass distribution toward higher MW PPSA assemblies (Supplementary Figure 3), we investigated the occurrence of heteromeric forms. Several proteins were consistently co-purified with PPSA during the sucrose gradient fractionation, with one stronger band around 70kDa (Supplementary Figure 12). Densitometry measurements on consecutive fractions separated by SDS-PAGE showed a sub-stoichiometric ratio of approximately 1:20 in different cell batches and purification rounds (Supplementary Figure 13), suggesting one copy per PPSA complex. The identity of this interacting protein was obtained from 4 crosslinks detected in 6 independent replicates (3 for SG and 3 for SEC) by XL-MS. This indicates that Phosphoenolpyruvate carboxykinase (PCK, approximately 70kDa) stably interacts with PPSA, as it survives extensive fractionation procedures prior to the crosslinking step. To further strengthen these findings, a co-evolutionary analysis was performed at the sequence level(Hopf et al., 2019), which uncovered 8 evolutionarily coupled sites at *>*99% confidence that mainly reside in the CD domain of PPSA (Supplementary Data 6). These coupled pairs combined with the distance constraints derived from XL-MS were further refined using DisVis to eliminate possible false positives(van Zundert et al., 2017), ultimately resulting in a refined set of 6 restraints (4 crosslinks; 2 evolutionarily coupled sites) that were used to dock PCK to the PPSA tetramer in its 3 main conformations: the “resting” state and CD-PBD or CD-NBD functionally active states (see Supplementary Note 1 and Supplementary Figure 13). Its position was further confirmed by the presence of a low resolution EM density (Supplementary Figure 13). Interestingly, PCK-PPSA complex formation is not possible in the CD-PBD state implying there is a specific functional role of PCK in locally hindering PPSA’s CD structural dynamics, thus PPSA catalysis (Supplementary Figure 13).

## Discussion

Metabolic enzymes are fundamental to life and as such evolutionarily, and thus structurally, constrained to maintain similar catalytic cores across species. Driving this maintenance, protein domains responsible for interactions between subunits or other proteins as well as the catalytic domains need to harbor certain structural features - especially for enzymes requiring large conformational changes such as the PPSA/PPDK class of enzymes(Kap Lim et al., 2007). We here thoroughly characterize an archaeal PPSA that has many structural similarities with the most studied PPDK-like enzymes in plants. These similarities strikingly already start from the conserved phosphorylatable sites THR-440 and SER-441(Chen et al., 2014; Reiland et al., 2009) of the CD mobile domain (Figure 4A), the building block for converting PEP to pyruvate. We uncover the phosphorylation site as Thr440 based on our quantitative and qualitative assessment of the proteomics data we recorded. Besides phosphorylation we also uncovered a plethora of lysine acylations, most of them understudied. Among these, acetylation is here the most abundant in multiprotein complexes, a PTM that was previously reported to be approximately 5 to 10 times more abundant than phosphorylation in yeast protein complexes(Šoštarić et al., 2018) and chloroplast ATP synthase(Schmidt et al., 2017). This class of PTMs fulfill a structurally stabilizing role at protein interfaces inside the ATP synthase complex for highly mobile proteins (Schmidt et al., 2017), where their placement is enzymatically driven. In our study we found acetylated lysines widely spread in *P. furiosus* protein complexes, not only at interaction interfaces but more generally in solvent accessible residues. In addition, we did not observed any clear sequence motifs supporting enzymatic acetylation or de-acetylation, suggesting an ulterior route for lysine acetylation. It was previously reported that cysteine residues in close structural proximity within proteins that reside in eukaryotic mitochondria can act as a carrier for non-enzymatic transfer of the acetyl group from acetyl-CoA(Wagner and Payne, 2013) or acetyl-phosphate. Such “accessory” cysteine residues not engaged in the catalytic pocket or in forming stabilizing disulfide bonds are indeed present in the PBD and CD domains of PPSA (CYS-746, 454 Supplementary Figure 11). By utilizing state-of-the-art MD simulations, we were able to demonstrate how PPSA structural dynamics and organization adapted to the occurrence of these environmental chemical modifications. Ultimately, only in the presence of acetylated lysine residues the outer NBD domains and their inter-domain interfaces are structurally stable at elevated temperatures. Strikingly, catalytically relevant CD-PBD contacts can only be sampled when acetylations are present in our MD simulations, further underlying a specific functional role for catalysis. Remarkable is that we could simulate this functional state with coarse grained modeling of a 2.15 MDa protein complex without including cofactors, thus paving the way for analyzing the structural effects of PTMs on large and flexible structures in silico. The 12 independent trajectories of the tetramers also highlighted the variability within the same arrangement, even if starting from the same model and the same (non-)acetylation pattern. When placing this in a cellular context, with the wide range of spontaneous lysine acylations detected in this work, the relatively slow kinetics this enzyme exhibits in vitro (approximately 20 ms turnover) is not surprising, ultimately explaining the massive abundance of this pivotal enzyme for the central carbon metabolism. Our findings additionally have ramifications on an evolutionary point of view. Non-enzymatic acetylation was found to occur extensively in the “prokaryotic-like” mitochondrial compartments of mammals (Wagner and Payne 2013), and tens of thousands of “off target” sites not linked to specific acetyltransferase have been reported that account for as high as 63% of mitochondrial proteins in mice (Baeza 2016). Given the compartmentalization of the cellular processes in Eukaryotes, it is likely that non-enzymatic acetylation became obsolete in mesophilic environments where the pH, temperature and availability of acetyl-containing chemical intermediates, or a combination thereof, became less favorable. As proteins evolved their functions with acetylation as a key component, an ulterior strategy was required for tuning these functions and replacement by an enzymatic-dependent process has been the likely outcome.

Large multimeric assemblies of archaeal PPSA from Staphylothermus marinus were the focus of several structural studies of two decades ago (Cicicopol et al., 1994, 1999; Harauz et al., 1996; Li et al., 2000). These were limited by the state of the technology and the intrinsic flexibility and conformational heterogeneity of the outer CD and NBD domains. Here, we confirm most of the previously reported findings, including the C-terminal α-helical oligomerization motif, the ancient origin of this motif, and its conservation – even in some hyperthermophilic sulphate-reducing bacteria (Supplementary Figure 8). The coordination of a Fe(II) atom restricts the structural organization to primordial anaerobic environments where ferrous iron was abundant until the oxygenation of the atmosphere led to its oxidation(Ilbert and Bonnefoy, 2013). We extend on the previous observations by showing how the tetrameric module revolving around the α-helical oligomerization motif of the PBD is the minimal functional PPSA unit, retaining catalytic activity upon disruption of the 24-mer oligomer (Supplementary Figure 3). This minimal unit is however not stable at physiologically relevant temperatures (above 80°C) and cytosolic concentrations, creating a strong evolutionary pressure for the emergence of the 24-meric oligomer. Interestingly, the dimeric interfaces between PPSA monomers of neighboring tetrameric modules are a conserved feature in bacterial and eukaryotic PPDK dimeric arrangements (Supplementary Figure 7). Although it is tempting to conclude that the large oligomeric organization of PPS in the two (unrelated) hyperthermophilic Archaea *P. furiosus* and *S. marinus* is an adaptation to increase its thermo stability, further research will be needed. A good starting point would be to investigate the PPS proteins with a motif predicting a 24-meric assembly found in mesophilic bacteria, and those without such a motif found in hyperthermophilic Archaea (as shown in Supplementary Figure 7). PPSA represents roughly 2% (this work) to 5%(Hutchins et al., 2001) of the cytosolic protein content, which is rather unusual for a metabolic enzyme as cells tend to produce these proteins in the minimal amount needed to support a given metabolic flux and economize energetic resources(Noor et al., 2016). The massive accumulation of PPSA appears to be linked to a low thermodynamic driving force combined with a relatively slow rate due to the inefficiencies of the swiveling domain mechanism at the extreme temperatures this organism thrives in. Besides increased thermostability, other explanations are also conceivable, *e*.*g*., it might somehow facilitate the (unusual) glycolytic role PPSA has in *P. furiosus*, opposed to the typical gluconeogenic role. Additionally, the high abundance of PPSA – presumably the result of its slow kinetics and inefficient (glycolytic) thermodynamics – also creates a strong evolutionary drive to adopt moonlighting functions, *i*.*e*. the adoption of secondary functions, something almost ubiquitously seen in abundant proteins such as glycolytic and ribosomal proteins(Copley, 2014). Glyceraldehyde 3-phosphate dehydrogenase, the quintessential moonlighting protein with dozens of secondary functions ascribed to it in Prokaryotes and Eukaryotes(Sirover, 2016), is also notoriously slow and typically one of the most abundant proteins. The uncovered interaction between PPSA and PCK potentially fits moonlighting. Its association hinders PPSA’s CD domain structural dynamics and thus catalysis, likely triggering a local switch between metabolic pathways depending on availability of ATP/AMP (PPSA substrate) or GTP (PCK substrate). This, in fact, might already be a first hint towards a possible moonlighting function stemming from PPSA’s cellular abundance and large oligomeric organization.

## Conclusions

In this study we characterized an intrinsically flexible and heterogeneous mega-dalton sized enzyme using an integrative structural approach that exceeds the sum of the parts of the state-of-the-art technologies. The structural insights into the large PPSA 24-mer, challenging common assumptions of required rigidity of thermostable proteins, are of prime interest for protein design approaches(Divine et al., 2021; Hsia et al., 2021) to enhance our ability to construct proteins that mimic its functional stability. From an evolutionary point of view, the structural and sequence similarities among eukaryotic bacterial, euryarchaeal and crenarchaeotal arrangements suggests an ancient origin of this hyperthermoactive PPSA 24-mer. Finally, non-enzymatic lysine acetylation is a structurally stabilizing PTM in the context of the flexibility required for the structural rearrangement of PPSA during catalysis. This PTM, along with other chemically driven acylations, might therefore also be of archaic origin, considering that acetyl-CoA (likely the substrate for non-enzymatic acetylation) may form pyruvate from H_2_ and CO_2_ without enzymes under hydrothermal vent conditions – relevant for tracing back the earliest life form(Martin, 2020; Weiss et al., 2016). The excessive presence of chemically modified lysines on protein surfaces, enhanced by the extreme environmental conditions, represents a paradigm shift in our understanding of the origin and role of PTMs in protein structural dynamics. Shifting from an environmentally occurring reaction, proteins had to cope with in the earliest, non-compartmentalized, organisms and in prokaryotic-like organelles like mitochondria, to a precise enzyme mediated, on/off function in specific cellular processes of eukaryotes.

## Methods

### Purification of PEP synthase from *P. furiosus*

*P. furiosus* cells (DSM 3638) were obtained from the Archaea Centre Regensburg (Institute of Microbiology and Archaea Centre at the University of Regensburg). Cells were grown at 95 °C in a 300 L fermenter on SME medium (Blöchl et al., 1997) containing 0.1 % (w/v) starch, 0.1 % (w/v) yeast extract (YE) and 0.1 % (w/v) peptone. Cell pellets were stored at −80 °C prior to lysis. For each lysis round, 1.5 g of cells were resuspended in 10 mL of lysis buffer (LB, 25 mM HEPES pH 7.4, 150 mM NaCl, 2 mM MgCl_2_) supplemented with 0.3 mg/mL DNAse I (Roche), 1 tablet of phosphatase inhibitor (PhosSTOP, Roche) and 1 tablet of protease inhibitor (EDTA free, Roche). Resuspended cells were subjected to Nitrogen cavitation (Simpson, 2010) in a pressurized vessel at 250 bar for 10 min. The lysate was fractionated by sequential centrifugation at 1500 g for 10 min (P1), and the supernatant was further centrifuged at 20,000 g for 20 minutes (P2) and subsequently at 100000 g for 60 minutes (P3). All lysis and centrifugation steps were conducted at 4 °C or on ice. The protein concentration of each fraction was evaluated with a BCA protein assay (Pierce, Thermo Scientific). For sucrose gradient ultracentrifugation, 450 µL of the final supernatant (S3, 12-13 mg/mL of protein) was loaded onto a tube with a 11 mL linear sucrose gradient (0.6 M Sucrose, 25 mM HEPES pH 7.4, 150 mM NaCl, 2 mM MgCl_2_). Gradient formation was done by freezing/thawing as previously described (Albanese et al., 2019). Separation was achieved through ultracentrifugation at 39,000 rpm for 16 h on a TH641 rotor (Sorvall, Thermo). Extraction of the fractions from the tube was done manually with a Hamilton 250 µL-syringe equipped with a custom-made needle reaching the bottom of the tube. As a second purification strategy, 500 µg clarified supernatant from P2 fractions was further fractionated by size-exclusion chromatography (SEC) on a Superose 6 increase column (GE Healthcare) equilibrated with LB buffer (25 mM HEPES pH 7.4, 150 mM NaCl, 2 mM MgCl_2_). SEC fractions were pooled according to their main elution peak (Supplementary Figure 1).

### Native and denaturing gel electrophoresis, densitometry quantification and in-gel digestion

Protein fractions and purified complexes were resolved by polyacrylamide gel electrophoresis (PAGE) in denaturing conditions after incubation into XT-sample buffer (Bio Rad) for 5 min at 90 °C with 12% Bis-Tris Criterion XT precast gels (Biorad, USA) with 1x XT-MOPS buffer at fixed current of 25 mA for 1.5-2 h. Proteins were stained in-gel using Imperial Protein Stain (Thermo Fisher Scientific). For non-denaturing Blue Native (BN) PAGE 10 µg of protein sample was mixed with 4x sample buffer (NuPAGE, Invitrogen) and subsequently loaded onto a Bis-Tris gel (3–12%). Electrophoresis was started with concentrated blue cathode buffer (1x) for 30 min at 70 V before the dark blue buffer was changed to a light blue cathode buffer (0.1x). After the change of buffer, the voltage was increased to 90 V for another 30 minutes, then 110 V for 1 h and 140 v for the last 1-2h. Gels were then fixed and stained with using Imperial Protein Stain (Thermo Fisher Scientific) and gel bands of interest excised and proteins digested as described earlier (Hevler et al., 2021).

### Mass Photometry

Mass photometry experiments to assess the oligomerization state under varying reducing conditions and temperatures were performed on a Refeyn OneMP instrument (Refeyn Ltd.). CultureWell gaskets (Grace Biolabs) were placed into the Microscope coverslips (24 mm × 50 mm; Paul Marienfeld GmbH). Three independent batches of purified PPSA at 0.1 mg/mL were diluted 700x in a buffer containing 0, 5, 10 or 15 mM Tris(2-carboxyethyl) phosphine hydrochloride (TCEP, Sigma-Aldrich). Samples in triplicates in the different reducing conditions were incubated at 20, 60, 80 or 90 °C for 10 minutes in a ProFlex PCR System thermal cycle (Applied Biosystems, Thermo Fisher Scientific) in MicroAmp Optical 8-Cap PCR tubes (Applied Biosystems, Thermo Fisher Scientific) to avoid evaporation. As each MP measurement takes approximately 1 minute, each replicate was independently incubated prior to the measurements. For each measurement, 13 µL of LB buffer was first placed into the well to adjust the focus, after that 2 µL of sample was mixed in. Movies were recorded for 60 seconds at 100 frames per second. A calibration measurement under the same conditions was performed roughly every 15 measurements using an in-house prepared protein standard mixture: IgG4Δhinge-L368A (73 kDa), IgG1-Campath (149 kDa), apoferritin (479 kDa), and GroEL (800 kDa). Data were processed using DiscoverMP (Refeyn Ltd.).

### Chemical crosslinking and bottom-up proteomics sample preparation, mass spectrometry acquisition and data analysis

For XL-MS sample preparation, 400 µg of Lysates (Supernatant of P1 centrifuge as described above) and 150 µg of purified PPSA from SG and SEC fractions at 1 mg/mL total protein concentration (measured by a Pierce BCA protein assay, Thermo Scientific, according to manufacturer instructions) were incubated with the crosslinking reagent PhoX(Steigenberger et al., 2019) for 30 min at room temperature. All replicates, unless specified, were purified from independent cell lysis batches from the two different batches of cells, as described above. The crosslinking reaction was quenched by addition of 1 M Tris·HCl to a final concentration of 10 mM. Crosslinked proteins were denatured in 8 M urea 50 mM Tris and sonicated for 15 cycles of 30 s with a Bioruptor Plus (Diagenode, SA) at 4°C. Disulfide bonds were reduced by addition of 8 mM DTT for 30 min at 54 °C, followed by alkylation with 16 mM IAA for 1 hour at room temperature in the dark. After dilution of the urea concentration to *<*2M with 50 mM Tris·HCl, proteins were digested with a combination of LysC (Wako Chemicals) and Trypsin (Porcine, Sigma Aldrich) (1:70 and 1:25 enzyme to protein ratio, respectively) for 16 h at 37 °C. Trifluoroacetic acid (TFA) at a final concentration of 0.5% was added to quench the digestion. Prior to Fe-IMAC enrichment, peptides were desalted by HLB Oasis (Waters) according to manufacturer instructions with the exception of using 0.1% TFA to get pH 2 in all buffers. The desalted peptides were lyophilized by vacuum centrifugation to near-complete dryness. Crosslinked peptides were enriched with Fe(III)-NTA cartridges (Agilent Technologies) using the Bravo AssayMAP robot (Agilent Technologies) as previously described (Steigenberger et al., 2019) and peptide mixtures were lyophilized by vacuum centrifugation to near-complete dryness and stored at −20 °C.The final peptide mixture was resuspended in 2% (v/v) formic acid prior to LC-MS/MS data acquisition.

Mass Spectrometry data were acquired using two UHPLC systems, an Agilent 1290 system or Ultimate 3000 RSLC nano system (Thermo Scientific) coupled to different models of Orbitrap mass spectrometers (Thermo Scientific), namely Exploris 480, Q-Exactive HF-X or Fusion tribrid. The MS setup of each raw file is detailed in the Supplementary Table 4. The experimental design was conceived to ensure maximum reproducibility within the same batch of samples. Peptides are first trapped in a pre-column (Dr. Maisch Reprosil C18, 3 µm, 2 cm × 100 µm) prior to separation on the analytical column packed in-house (Poroshell EC-C18, 2.7 µm, 50 cm × 75 µm), for samples run on the Ultimate 3000 RSLC nano system setup both columns were kept at 40 °C in the built-in oven. Trapping was performed for 10 min in solvent A (0.1% v/v formic acid in water), and the elution gradient profile was as follows: 0 – 10% solvent B (0.1% v/v formic acid in 80% v/v ACN) over 5 min, 12 - 37% solvent B over 55 min, 40-100% solvent B over 3 min, and finally 100% B for 4 min. The mass spectrometer was operated in a data-dependent mode. Full-scan MS spectra were collected in a mass range of m/z 350 – 1300 Th in the Orbitrap at a resolution of 60,000 at m/z=200 Th after accumulation to an AGC target value of 1e6 with a maximum injection time of 50 ms. In-source fragmentation was activated and set to 15 eV. The cycle time for the acquisition of MS/MS fragmentation scans was set to 3 s. Charge states accepted for MS/MS fragmentation were set to 3 - 8. Dynamic exclusion properties were set to n = 1 and to an exclusion duration of 15 s. Stepped HCD fragmentation (MS/MS) was performed with increasing normalized collision energy (19, 27, 35 %) and the mass spectra acquired in the Orbitrap at a resolution of 30,000 at m/z=200 Th after accumulation to an AGC target value of 1e5 with an isolation window of m/z = 1.4 Th and maximum injection time of 120 ms.

The acquired raw data were processed using Proteome Discoverer (version 2.5.0.400) with the integrated third-party XlinkX/PD nodes. Normal peptide search was performed using the Mascot node against the full *P. furiosus* UniProtKB database (2566 entries on 09/02/2020) in silico digested with Trypsin/P with a minimal peptide length of six and two miss cleaved sites allowed. Cysteine carbamidomethylation was set as fixed modification. Methionine oxidation, protein N-term acetylation and lysine acetylation were set as dynamic modifications. For the search of mono-linked peptides, water-quenched (C_8_H_5_O_6_P) and Tris-quenched (C_12_H_14_O_8_PN) were set as dynamic modifications on lysine residues. Filtering at 1% false discovery rate (FDR) at the peptide level was applied through the Percolator node. For crosslinked peptides, a database search was performed using XlinkX/PD nodes against a database containing the most abundant 100 proteins identified from the normal peptide search in the Mascot node, with the same protease and dynamic modifications allowed but an increased number of missed cleavages of 3. Identified crosslinks were only accepted through the Validator node at 1% FDR with a minimal score of 40 and a minimal delta score of 4.

For bottom-up proteomics measurements, 50 µg of protein material was prepared as described above, without incubation with the crosslinking reagent and without enrichment prior to MS analysis. Differences for the acquisition times applied for full lysate measurements where the gradient was as follows: 9-13% solvent B (0.1% v/v formic acid in 80% v/v ACN) over 5 min, 13-44% solvent B over 95 min, 44-99% solvent B over 2 min, and finally 99% B for 4 min (flow was set to 300 nL/min) prior to re-equilibration with solvent A for 5 minutes. Mass spectrometry data was collected in a data-dependent fashion with survey scans from m/z 300 to 1500 Th (resolution of 60,000 at m/z=200 Th), and up to 15 of the top precursors selected for fragmentation (MS/MS) using a normalized HCD collision energy of 27%. The MS2 spectra were recorded at a resolution of 30,000 (at m/z=200 Th). The AGC targets for both MS and MS2 scans were set to standard with a maximum injection time of 50 and 35 ms, respectively. Raw data were processed using the MaxQuant software(Cox and Mann, 2008) version 1.6.17 with standard settings applied. Briefly, the extracted peak lists were searched against the reviewed Human UniProtKB database (date 15-07-2021; 20353 entries), with a precursor mass deviation of 4.5 ppm and a fragment mass deviation of 20 ppm. Cysteine carbamidomethylation was set as static modification, and methionine oxidation, N-terminal acetylation, lysine acetylation and serine/threonine/tyrosine/histidine phosphorylation as variable modifications (maximum 5 modifications per peptide allowed). For the acylation search the following modifications were set on non-terminal lysine residues (UNIMOD accession code in brackets): Propionylation (58), Gluratylation (1848), Crotonylation (1363), Butyrylation (1289), Malonylation (747). Both LFQ quantification and “match between runs” were enabled. Lysine acylation and phosphorylation stoichiometry for each site was calculated by the MaxQuant software as a mod/base ratio (intensity of the modified peptides vs. the intensity of its unmodified counterpart). Additional spectral inspection and localization probability calculations were performed on each fragmentation spectrum using the in-house software FragmentLab v2.6.1.7 (available for download at https://scheltemalab.com/software). Only phosphorylated sites with a localization probability of 1 and mod/base ration of 0.5 were further considered. Of the 51 acetylated lysines found for PPSA, a subset of 14 “highly acetylated lysine” residues were selected if found in all samples types (in-gel digested BN bands and SG purified complexes) and the occupancy was *>*10% (Supplementary Figure 10). The iBAQ values in Figure 1A are approximate absolute abundances of the identified proteins derived by the normalization of the summed peptide intensities by the number of theoretically observable peptides for a given protein (Schwanhüusser et al., 2011).

### Cryo-EM Single Particle analysis data acquisition and image processing

Fractions of 100 µL were collected from 6 sucrose gradient tubes and each fraction checked for purity by SDS-PAGE. Cleaner fractions were pooled, concentrated on a 100 kDa cellulose cut-off filter (5 mL, Millipore) centrifuging at 2500 g for 15 minutes at 4 °C. Sucrose was removed by buffer exchanging in LB buffer and further concentrated to 2 mg/mL prior to cryo-grid preparation. 3 µL of purified PPSA was applied to glow-discharged holy grids (Quantifoil, Cu200, R2/2) for 30s. The grids were blotted for 3.5s then plunged into liquid ethane using an FEI Vitrobot Mark IV (Thermo Fisher Scientifc) at 100% humidity and 25 °C. PPSA cryo-EM data were collected using a Titan Krios microscope (Thermo Fisher Scientific) operating at an accelerating voltage of 300 keV. All datasets were automatically recorded at a defocus range of −0.5 µm to −2 µm with a pixel size of 0.66 by the EPU software (Thermo Fisher Scientific). A total of 1842 images were collected on a Falcon camera in electron counting mode. Each image was dose-fractionated into 40 frames with a total dose of 40 electrons per Å ^2^.

For the PPSA PBD domain structure determination, all image processing was performed using RELION 3.1(Zivanov et al., 2018). All images were motion corrected and electron-dose weighted with RELION’s MotionCor2 algorithm (Zheng et al., 2017)(Zivanov et al., 2018). CTF estimation was determined with RELION’s CTFFIND-4.1 program(Rohou and Grigorieff, 2015). Images whose estimated resolution from CTFFIND-4.1 were worse than 5Å were discarded, leaving 1171 images for further data processing. Using RELION’s Laplacian picker, 96782 particles were automatically picked and binned, and then subjected to 2D classification. After 1 round of 2D classification, particles from classes not showing clear secondary structural features were removed. A total of 18954 selected particles were subjected for the initial model creation with octahedral symmetry imposed, followed by several rounds of auto-refinement and 3D classification and auto-refinement, yielding an estimate resolution of 3.77 Å. Five runs of CTF-refinement (optical aberration correction, magnification anisotropy, per-particle defocus and per-micrograph astigmatism) yielding a final resolution of 2.90 Å.

To obtain the density map of the flexible CD domain, another 8,000 images were collected with a pixel size of 0.83 Å on a Titan Krios microscope (Thermo Fisher Scientific). Images for which the estimated resolution was worse than 5 Å were removed, leaving 6,607 images for further data processing. Using RELION’s Laplacian picker, 578,504 particles were automatically picked and binned to 1.66 Å per pixel, and then subjected to 2D classification. After 1 round of 2D classification, 250,258 particles were used for the initial model creation with octahedral symmetry imposed, followed by several rounds of auto-refinement and 3D classification and auto-refinement, yielding an approximate resolution of 3.4 Å. Octahedron symmetry expansion was applied to this 3.4 Å density map and focused classification performed in RELION 3.1.1. To do so, all the particles in the final auto-refinement were duplicated 24 times and the local refined density map was used as a reference for 3D classification without particle alignment using a mask solely enclosing the protein portion. The best class was reconstructed to a final resolution of 7.1Å. All the image processing procedures were summarized in Supplementary Figure 4.

### Model building and refinement

The initial model was predicted by AlphaFold and fitted into the cryo-EM density map using UCSF Chimera (Pettersen et al., 2004). The poorly built model was removed, several rounds of refinement were applied by Phenix with secondary structure restraints and geometry restraints, followed by manual adjustments of the model with Coot 0.9.8 (Emsley and Cowtan, 2004). The CD domain model was extracted from the full length prediction with the trROSETTA webserver (accessed at: https://yanglab.nankai.edu.cn/trRosetta/) (Yang et al., 2020) and docked into the EM density map using UCSF Chimera (Supplementary Figure 4). The map-model FSCs were calculated using PHENIX Mtriage (Afonine et al., 2018), with model statistics provide in Supplementary Table 6. All figures were generated with UCSF ChimeraX (Goddard et al., 2018).

### Structural modelling of three PPSA tetrameric states

In its catalytic cycle, PPSA (Uniprot code P42850) samples a variety of conformations and performs two reactions. In the first one, a phosphate group from ATP is transferred to CD at the NBD active site, phosphorylating a CD residue (CD-NBD state). This is followed by a conformational change that relocates the phosphorylated CD domain to the PEP active site on PBD (CD-PBD state). Then the second reaction is performed, transferring the phosphate group to a pyruvate substrate, yielding phosphoenolpyruvate. Due to the large conformational changes that PPSA undergoes during PEP production, three proposed PPSA states corresponding to different steps in the catalytic cycle have been modelled. These three states represent conformations in which the central domain is interacting with i) the active site of the Nucleotide Binding Domain (CD-NBD state), ii) the active site of the Pyruvate Binding Domain (CD-PBD state) and iii) with both domains (NBD and PBD) where the CD is not located near either of the two active sites (resting state).

The complete PPSA oligomer is composed of 24 units (6*4 tetramers), therefore, each PPSA state was modeled in its homo tetrameric form. The PPSA PBD region was resolved in the cryo-EM data (see Cryo-electron microscopy and single particle analysis section). In the setup of all three states, the cryo-EM PBD structure (residues 507-817) as well as its orientation within the tetramer conformation of the PPSA 24-mer were maintained. NBD (residues 1-360) and CD (residues 380-482), however, were not resolved at sufficient resolution in the obtained cryo-EM data to resolve their structures at atomic level. Therefore, they were modelled with the trROSETTA webserver (accessed at: https://yanglab.nankai.edu.cn/trRosetta/) (Yang et al., 2020), for the residues 1-484. The structure of the two domains combined and their orientation with respect to PBD was modelled using different experimental and bioinformatics techniques as described below.

**The CD-NBD state** was predicted by:

1. **Predicting the intermolecular contacts between CD-NBD and PBD –** To predict the orientation of the CD-NBD region with respect to PBD, the co-evolution tools EVcomplex and RaptorX (Hopf et al. 2019; Zeng et al. 2018) were both applied to the PPSA system, using the CD-NBD (residues 1-483) and PBD regions (residues 514-817) to predict intermolecular contacts between the two regions.
2. **Filtering the predicted contact data –** The top 5 intermolecular contacts with the highest probability score from EVcomplex and RaptorX, were merged to prepare a residue-residue contact list for DisVis(van Zundert and Bonvin 2015; van Zundert et al. 2017). The resulting list of 10 contacts, together with the structures of the modelled CD-NBD and resolved PBD were used in DisVis to identify possible false positives. For DisVis, these contacts were defined by a Cα-Cα distance restraint with a maximum length of 10 Å. The DisVis analysis was run using the Complete Scanning option on the webserver (https://wenmr.science.uu.nl/disvis/submit) with a voxel spacing of 1 Å and a rotational sampling angle of 9.72°.
3. **CD-NBD and PBD docking with filtered contacts –** The DisVis-ranked contacts with a negative z-score, the contacts most likely to be at the protein-protein interface, were selected as unambiguous restraints for protein-protein docking with HADDOCK2.4(van Zundert et al. 2016). These contacts included Lys277-Glu570, Glu276-Lys574 and Asp95-Glu558 for which the maximum Cα-Cα distance was set to 10 Å and the lower bound threshold to 0 Å, in agreement with the DisVis setup. Using these three unambiguous restraints, models of the interaction between the CD-NBD region (residues 1-484) and PBD region (residues 507-817) were obtained with the HADDOCK2.4 webserver (https://wenmr.science.uu.nl/haddock2.4/). A total of 4 clusters were predicted, which were analysed to identify a model that would not structurally overlap with the tetramer and the 24-mer PPSA conformations of the cryo-EM resolved PBD.
4. **Modelling the CD-PBD linker –** The next step was to model the missing linker between CD and PBD to obtain a complete PPSA monomer. This linker consists of 31 residues (residues 483-513), for which no structural templates are available. PEP-FOLD3 (Lamiable et al., 2016) was used to predict the 3D conformation of the linker.
5. **Modeling CD-NBD PPSA state –** The predicted helical linker region (residues 484-499) was combined with the docked CD-NBD:PBD conformation in a PPSA octamer form (extracted from the PPSA 24-mer) as a template for MODELLER version 9.12(Eswar et al., 2006) to predict the complete PPSA monomer in line with the PPSA 24-mer system. A monomer of this modeled PPSA state was aligned with the cryo-EM PBD data to set up a second MODELLER run in which residues 499-507 were remodeled, leading to a symmetric PPSA tetramer with all monomers in the CD-NBD state. The experimentally identified Cys42-Cys189 disulfide bond was included as well.A total of 60 models were generated per MODELLER run and ranked by their DOPE score. The top-10 ranked models were used for model selection. The highest ranked model of the top 10 was selected for structure refinement when loop entanglement was absent (i) and the lowest RMSD in the PBD region of the tetramer was obtained with respect to the template (ii).
6. **Including acetylated lysines in the modeled CD-NBD state of PPSA –** The predicted PPSA tetramer conformation was refined with HADDOCK2.4 to optimize the intermolecular protein-protein interfaces within the PPSA tetramer and to include the experimentally identified acetylated lysines (106, 120,185, 187, 427, 466, 492, 496, 557, 574, 641, 726, 737, 805). The residue name of those residues was changed to ALY in the PDB file provided to HADDOCK to indicate they had to be generated by HADDOCK as acetylated lysines.

**The CD-PBD state** was predicted by:

1. **Reorienting CD’s position on PBD –** While the NBD position with respect to PBD of the CD-NBD state was maintained, the CD region had to be relocated in the CD-PBD state as this conformation should position the temporarily phosphorylated His442, the equivalent residue to His456 in PPDK (Uniprot code P22221), in close proximity to the PEP active site on PBD to take part in the second PPSA reaction mechanism, pyruvate to PEP conversion. Therefore, CD (residues 379-483) was relocated according to a PPDK structure in a CD-PBD like conformation (PDB ID 5JVL, chain C). First the PBD of PPDK was aligned to the PBD of PPSA. Subsequently, an isolated PPSA CD was aligned on PPDK CD, leading to a PPSA CD region that interacts with the PPSA PEP active site on PBD. These orientations of NBD, CD and PBD were merged to form a PPSA template that was then used in MODELLER.
2. **Predicting the CD-PBD and CD-NBD linkers –** Due to this change in the CD position, the linkers between CD and PBD (CD-PBD linker) and CD and NBD (CD-NBD linker) had to be adjusted. Both linkers were run through PEP-FOLD3. For the CD-PBD linker, a helical element was identified for residues 484-499, while for the CD-NBD linker a helical element was predicted for residues 363-378.
3. **Modeling CD-PBD PPSA state –** The predicted helical linker regions (residues 363-378 and 484-499) were combined with the NBD-CD-PBD domain template to build a PPSA tetramer template (PDB regions aligned with the tetramer from the PPSA 24-mer) to model the complete form of the PPSA monomer in line with the PPSA 24-mer system. A monomer from this modeled PPSA state was aligned with the cryo-EM resolved PBD tetramer to set up another MODELLER run in which residues 499-507 were remodeled, resulting in a more symmetric PPSA tetramer with all monomers in the CD-PBD conformation. A total of 60 models were generated per MODELLER run and these were ranked by their DOPE score. The top-10 ranked models were used for model selection. The highest ranked model of the top 10 was selected for structure refinement when loop entanglement was absent (i) and the lowest RMSD in the PBD region of the tetramer was obtained with respect to the template (ii).
4. **Including acetylated lysines in the modeled CD-PBD state of PPSA –** The model was refined with HADDOCK, introducing the identified acetylated lysines as described above for the CD-NBD state.

**The resting state** was predicted by:

1. Reorienting CD’s position on PBD – According to one PPSA monomer in the obtained cryo-EM data of the PPSA 24-mer, CD can be associated to PBD in the resting state without interacting with the PEP active site. To prepare the PPSA template to model this state, the predicted CD-PBD state was used as a starting structure. While the location of the PBD, residues 508-817, was maintained, the CD-NBD region (residues 1-502) was reoriented according to the central-domain electron density found for the PPSA monomer, the resting template.
2. Modeling the PPSA resting state – This resting template of a PPSA monomer was used to build a PPSA tetramer conformation in line with the PPSA 24-mer structure to setup a modelling run with MODELLER. A total of 100 models were generated in this MODELLER run which were ranked by their DOPE score. The top-20 ranked models were used for model selection. The highest ranked model of the top 20 was selected for structure refinement when loop entanglement was absent and the lowest RMSD in the PBD region of the tetramer was obtained with respect to the template (ii).
3. Including acetylated lysines in the modeled resting state of PPSA – The model was refined with HADDOCK, introducing the identified acetylated lysines as described above. In addition, the phosphorylated form of Thr440 (defined as residue TOP for HADDOCK) was included for each monomer to mimic this experimentally confirmed phosphorylated state of PPSA. Another refined model was generated without the post-translationally modified lysines. These two resting state conditions, with and without acetylated lysines, were used as starting conformations for the coarse-grained molecular dynamics simulations described in the Molecular dynamics simulations and analysis section.

### Inductivity coupled plasma mass spectrometry (ICP-MS) elemental analysis

Precleaned polypropylene bottles were used for the preparation of all blanks, standards and samples. The bottles were rinsed with ultrapure water (18.2 MΩ·cm) and left to dry in a laminar flow clean hood before use. All blanks and calibration standards were prepared using 2% v/v HNO_3_ (68% Optima™ grade, Fisher Chemicals), with single element standards (Chloride is MERCK and others are SPEX CertiPrep). The samples were collected and made up 5mL with 2 v/v % HNO_3_. An internal standard solution containing Rh at 10 µg·L-1, was spiked into all samples for occurring matrix effects. An inductivity coupled plasma mass spectrometer iCAP TQe ICP-MS (Thermo Fisher Scientific, Bremen, Germany) was used for all measurements. The ICP-MS was operated in He KED mode using the parameters provided together with the quantification results and data in Supplementary Table 5.

### Phylogenetic and sequence motif analysis

PPSA homologous protein sequences were retrieved through NCBI-BLAST against the non-redundant protein sequences database. Four species per genus were selected among the top 2000 with a query coverage *>*75%. The resulting set of 325 sequences was aligned with Clustal Omega(Sievers et al., 2014) over 10 iterations and then manually curated using Gblocks(Talavera and Castresana, 2007). Maximum likelihood (ML) tree inference on the final set containing 978 distinct alignment patterns and 26% gaps was done with RAxML (version 8.2.12) testing three different substitution matrices (LG, JTT and GTR) over 100 bootstrap replicates. Sequence motif analysis for acetylated lysine residues was done with the IceLogo webserver(Ham et al., 2009) using the flanking 6 amino acids (+6 and −6) of the 51 acetylated lysines of PPSA (Supplementary Table 1). An analysis against a precompiled Swiss-Prot background database, a built-in function in the iceLogo webserver, produced similar results.

### PPSA activity assays

Catalytic activity was tested for both PEP- and pyruvate-forming reactions adapting the protocol described earlie (Hutchins et al., 2001). For the PEP-forming assay, the reaction mix contained 50 mM HEPES buffer (set to pH 8.84 at room temperature in a 0.5 M stock solution, corresponding to pH 8 at 80 °C), 20 mM KCl, 10 mM MgCl_2_, 20 mM NH4Cl, 4 mM sodium-pyruvate, 1.6 mM ATP, and 2.5-5 nM phosphoenolpyruvate synthase, in a volume of 1 ml. All 1.5-ml tubes with the reaction mix were incubated in a heating block at 80 °C, and the reactions were started by addition of ATP. Six 100 µl samples were taken over a period of up to 20 minutes, which were immediately pipetted into a 96-well plate that was placed on ice to terminate the reaction. The concentration of PO4 in the samples was analyzed using the Malachite Green Phosphate Assay Kit from Sigma-Aldrich. This was done in a new 96-well plate by mixing 72 µl water with 8 µl of the samples, and 20 µl of the reagent from the kit. A PO4 calibration curve was included on the same plate, from which the absorbance at 620 nm was measured. The rate of PO_4_ formation was used to calculate the turnover number of the enzyme, which had to be corrected for non-enzymatic PO_4_ release (typically around 0.07 µM/s), determined from the negative controls without phosphoenolpyruvate synthase added.

For the pyruvate-forming assay, the reaction mix contained 50 mM Na-PO4 buffer (set to pH 7.66 at room temperature in a 0.5 M stock solution, corresponding to pH 7.5 at 80 °C), 20 mM KCl, 2.5 mM MgCl_2_, 20 mM NH4Cl, 2 mM phosphoenolpyruvate, 2 mM AMP, 25 nM phosphoenolpyruvate synthase, in a volume of 1 ml. All 1.5-ml tubes with the reaction mix were incubated in a heating block at 80 °C, and the reactions were started by the addition of phosphoenolpyruvate. Six 100 µl samples were taken over a period of up to 20 minutes, which were immediately pipetted into a 96-well plate that was placed on ice, terminating the reaction. The concentration of pyruvate in the samples was analyzed using a mixture containing 20 U/ml lactate dehydrogenase (from rabbit) and 2 mM NADH, in water. This was done in a new 96-well plate by mixing 100 µl water with 50 µl of the samples, and 50 µl of the lactate dehydrogenase-NADH mix. NADH has a high absorbance at 340 nm, while NAD+ does not. Lactate dehydrogenase reduces pyruvate to lactate, through the oxidation of NADH to NAD+. A calibration curve of sodium-pyruvate was included on the same plate, from which the absorbance at 340 nm was measured. The rate of pyruvate formation was used to calculate the turnover number of the enzyme, which was corrected using the negative controls without phosphoenolpyruvate synthase added.

### Molecular dynamics simulations and analysis

The all-atom structures of the full c1(X4) 24-mer PPSA models, carrying highly acetylated sites at 14 positions (106, 120,185, 187, 427, 466, 492, 496, 557, 574, 641, 726, 737, 805) or unmodified lysines were coarse grained (CG), mapping their atoms to the SIRAH force field (ff), that uses a classical Hamiltonian common to most all-atom potentials to describe particle–particle interactions, and recently extended to support the PTMs most commonly found on proteins(Garay et al., 2020). Both starting structures contained a disulfide bond between Cys42-Cys189 as well as phosphorylation of Thr440. All starting structures, simulation boxes and parameters files used for the minimization, equilibration and production are provided (Supplementary Data 5). Simulation was conducted in GROMACS 2020.4(Hess et al., 2008), for which code can be found here: https://doi.org/10.5281/zenodo.5636522. The 24-aly system consisted of 94128 CG atoms for the protein representation, solvated with 179271 WT4 CG water beads (Garay et al., 2020). A neutral charge was achieved adding 6617 NaW (Na+) and 5633 ClW (Cl-), corresponding to a concentration of approximately 150 mM NaCl concentration. The 24-lys system was also prepared accordingly and consisted of 93456 CG atoms for the protein, 186690 WT4 CG water beads and neutralized with 6503 NaW (Na+) and 5855 ClW (Cl-). Solvation was done using the default radii of 0.105 nm for atoms not present in the VdW database (vdwradii.dat) and then removing the WT4 molecules within 0.3 nm from the solute. In all cases, eventual clashes were relaxed during the solute-restrained energy minimization. Due to the length of the loops connecting the 3 domains, an alternative configuration of the tetramers where the CD-NBD domains are sitting on top of the neighboring PPSA subunit is also possible (see Supplementary Note 1). This was named “alternative” (a) configuration, as opposed to the “original” (o), thus producing 4 starting models: a24-aly, o24-aly, a24-lys and o24-lys (Supplementary Data 5). All these configurations were subjected to a first round of production consisting of 125 ns in duplicate to define the stable configuration for further extension of the simulation time to 500 ns.

The main stages of the simulation can be summarized as follows:1) solvent and side-chain relaxation by 2 stages of 20’000 steps of energy minimization, imposing positional restraints of 1000 kJ mol-1 nm-2 on the whole protein (stage 1) and only on backbone beads (GN and GO, stage 2); 2) solvent NVT ensemble equilibration with a first stage where the temperature was slowly increased from 303K to 363K in 7 steps of 4 ns each, and a second stage to equilibrate the protein by gradually releasing the positional restraints from 1000 kJ mol-1 nm-2 on the backbone beads (GN and GO) to 100 kJ mol-1 nm-2 on the C-terminus to compensate for the missing stabilization effect of the Met799-Fe cluster; 3) production simulation of an additional 80 ns in NPT ensemble at 363 K and 1 bar imposing positional restraints of 100 kJ mol-1 nm-2 on the C-terminus. Nonbonded interactions were treated with a 1.2 nm cutoff and PME for long-range electrostatics. An integration time-step of 15 fs was used during MD production runs. The system pressure was controlled by the Parrinello-Rahman barostat (Parrinello and Rahman, 1981) with a coupling time of 4 ps. The positional restraints during the production simulation were necessary to maintain the overall system at the high energy of the particles at 363 K (90 °C), the temperature mimicking the near-optimum temperature for PPSA catalytic activity. All simulations were run in duplicate. Although the simulations do not accurately reflect the possible dynamics of the protein in native conditions, the computational challenges of simulating *>*350000 atoms for the protein only drove us to use a CG representation of the system. To analyze the trajectories, every functional module (tetramer) was extracted from the simulation of the full 24-mer for each replicate, resulting in two sets of 12 independent trajectories (6 for each 24-mer) either with and without acetylated Lysine residues. To produce morphed movies and analyze secondary structure and interfaces, the CG tetramers trajectories were back mapped to atomistic detail using SIRAH tools via VMD. A back mapped atomistic model was produced every 35.7 ns, and subjected to 100 cycles of energy minimization in AMBER, resulting in 14 atomistic models describing each trajectory of a given tetramer over the 500 ns of simulation, further interpolated in ChimeraX using the morph command to produce the atomistic representation of the dynamics (Supplementary Data 5). RMSD calculation and trajectory analysis were done with the MDanalysis suite(Michaud-Agrawal et al., 2011).

Each probability density in Figure 5B and 5C is calculated over two independent coarse-grained molecular dynamics simulations of the last 250 ns of production of the PPSA 24-mer in the acetylated and non-acetylated form, using a snapshot frequency of 5 ns. In Figure 5B, the RMSD of the Cα atoms (equivalent to backbone GC beads in CG MD) of the NBD region (residues 1-365) in the tetramer form was calculated with respect to their conformation in the tetramer PPSA resting state. The six trajectories of all six tetramers included in the simulated PPSA complex, resulting in 3 µs ([250ns*6]*2) of accumulated tetramer simulation, were used in the probability density calculation. In Figure 5C, the RMSD of the Cα atoms (equivalent to backbone GC beads in CG MD) was calculated over the CD (residues 379-481) and PBD (residues 510-790) regions with respect to the modelled CD-PBD state of PPSA in the monomer form. All 24 PPSA monomers of which the simulated PPSA complex was composed were included in the probability density calculation, resulting in 12 µs ([250ns*24]*2) of accumulated PPSA monomer simulation.

## Acknowledgments

This work is part of the research programme NWO TA with project number 741.018.201, financed by the Dutch Research Council (NWO). We further acknowledge funding through the European Union Horizon 2020 program INFRAIA project Epic-XS (Project 823839) and support of the and the Netherlands Electron Microscopy Infrastructure. PA acknowledge the Dutch National Supercomputer, supported by NWO, for the computational resources (grant agreement EINF-894).

## Contributions

PA – experimental design, conceptualization, protein purification, (Crosslinking) Mass Spectrometry, Mass photometry data acquisition and analysis, biochemistry, MD simulations and analysis, structural modelling, phylogeny, data analysis, data interpretation, manuscript writing. WS – Cryo-EM data collection, model building, manuscript writing. SvK – computer aided protein model building, interpretation of the molecular dynamics simulations, manuscript writing. JK – cell culture, activity assays, biochemistry, manuscript writing. FK – Cryo-EM data collection. BS - (Crosslinking) Mass Spectrometry, synthesis of reagents. TV – ICP-MS experiments. AK – funding, management. AJRH – funding. SWMK – funding, management, data interpretation. AMJJB – funding, management, data interpretation. FF – funding management, data interpretation. RAS – funding, management, experimental design, conceptualization, data analysis software, data interpretation, manuscript writing.

## Data availability

Atomic coordinates of the PPSA core are deposited in the PDB under accession code 8A8E. The EM density map has been deposited to the Electron Microscopy Data Bank under the accession EMD-15230. All MS raw data files are deposited to the ProteomeXchange Consortium via the PRIDE partner repository https://www.ebi.ac.uk/pride/archive/projects/PXD035847). The data to reproduce accurately the MD simulations (starting all-atom PDBs, CG models, GROMACS parameters, force field and topologies) are shared via the Zenodo repository and can be downloaded (https://doi.org/10.5281/zenodo.6961827). All other data needed to evaluate the conclusions in the paper are present in the paper and/or the Supplementary Materials.

