## Supplementary Table 6 for "Thriving in the heat – Lysine acetylation stabilizes the quaternary structure of a Mega-Dalton hyperthermoactive PEP-synthase"

Table | Cryo-EM data collection, refinement and validation statistics

|  | Composite map of PPSA core domain(EMD-15230, PDB 8A8E) |
| --- | --- |
| **Data collection and processing** | Titan Krios |
| Microscope | Falcon 4 |
| Voltage | 300 |
| Electron exposure(e^-^/ Å^2^ ) | 40 |
| Defocus range(um) | -1.0 to -2.0 |
| Pixel size (Å/pixel) | 0.88 |
| **Map resolution (Å)** | 2.9 |
| FSC threshold | 0.143 |
| **Model composition** |  |
| Chains | 24 |
| Atoms | 56832 |
| Residues | Protein: 7272 Nuleotide:0 |
| **Validation** |  |
| MolProbity score | 2.00 |
| Clashscore | 4.32 |
| Rotamer outlayers(%) | 2.81 |
| Ramachandran plot |  |
| Favored(%) | 93.00 |
| Allowed(%) | 7.00 |
| Disallowed(%) | 0 |
| **Refinement** |  |
| Correslation coefficient (CC mask) | 0.83 |
| Root-mean-square deviation (bond length)(Å) | 0.005 |
| Root-mean-square deviation (bond angles)(°) | 0.790 |
