## Supplementary material for "Thriving in the heat – Lysine acetylation stabilizes the quaternary structure of a Mega-Dalton hyperthermoactive PEP-synthase"

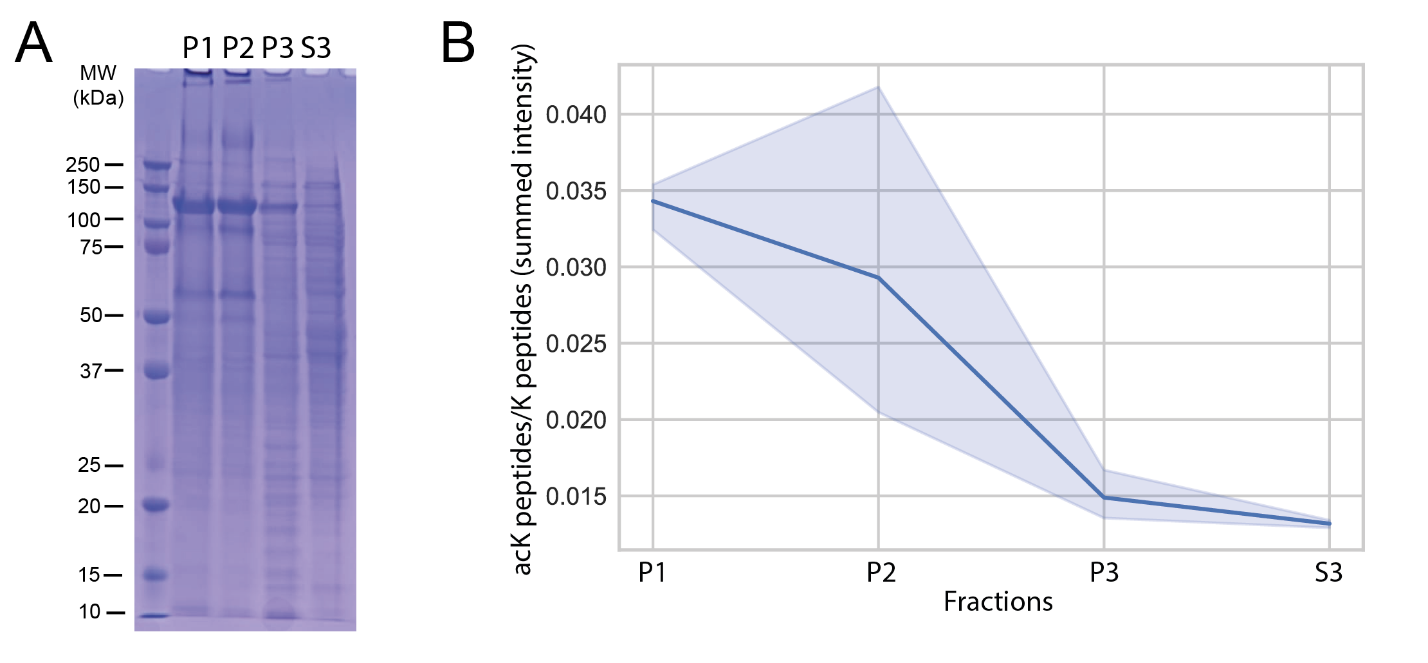
**Supplementary Figure 1. *P. furiosus* proteome fractionation by sequential centrifugation.** Cells lysed in mild conditions are fractionated and checked on SDS-PAGE **(A)**. The relative content of peptides containing acetylated lysine (acK) with respect to peptide containing unmodified lysine (*i.e.* K peptides) is plotted for 3 replicates for each sub-cellular fraction **(B)**.

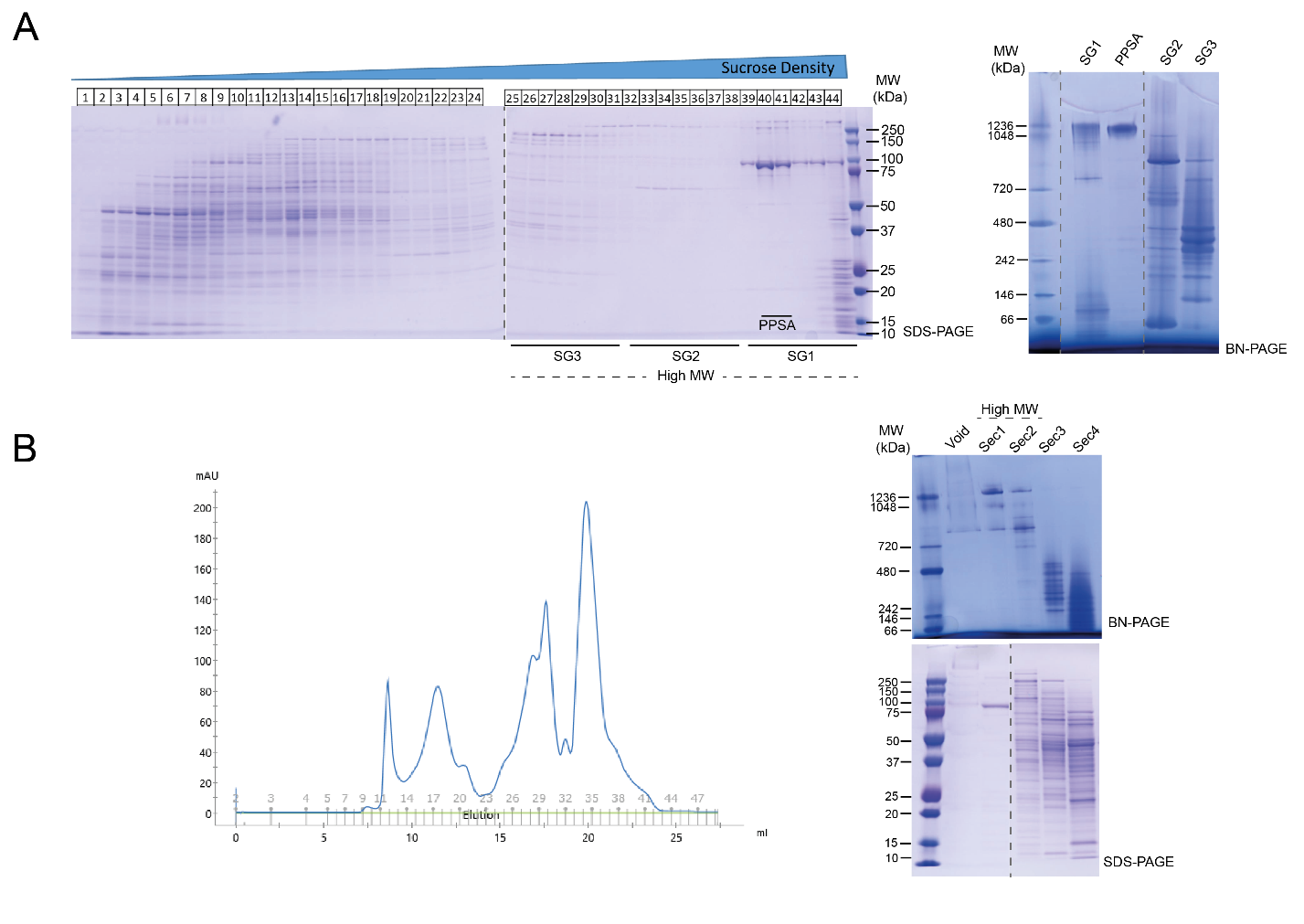

**Supplementary Figure 2. Native purification of PPSA through complete fractionation of the soluble cytosolic compartment.** Fraction S3 containing the cytosolic proteins was further separated by Sucrose gradient Ultracentrifugation (SG) **(A)** and Size exclusion Chromatography (SEC) **(B)**. Each fraction is then pooled in three main aggregated samples named SG- or SEC-1, 2 and 3 in decreasing order of molecular weight (MW). Each fraction or aggregated samples was checked by SDS- and BN-PAGE. The cleanest fractions containing PPSA subunits were collected separately.

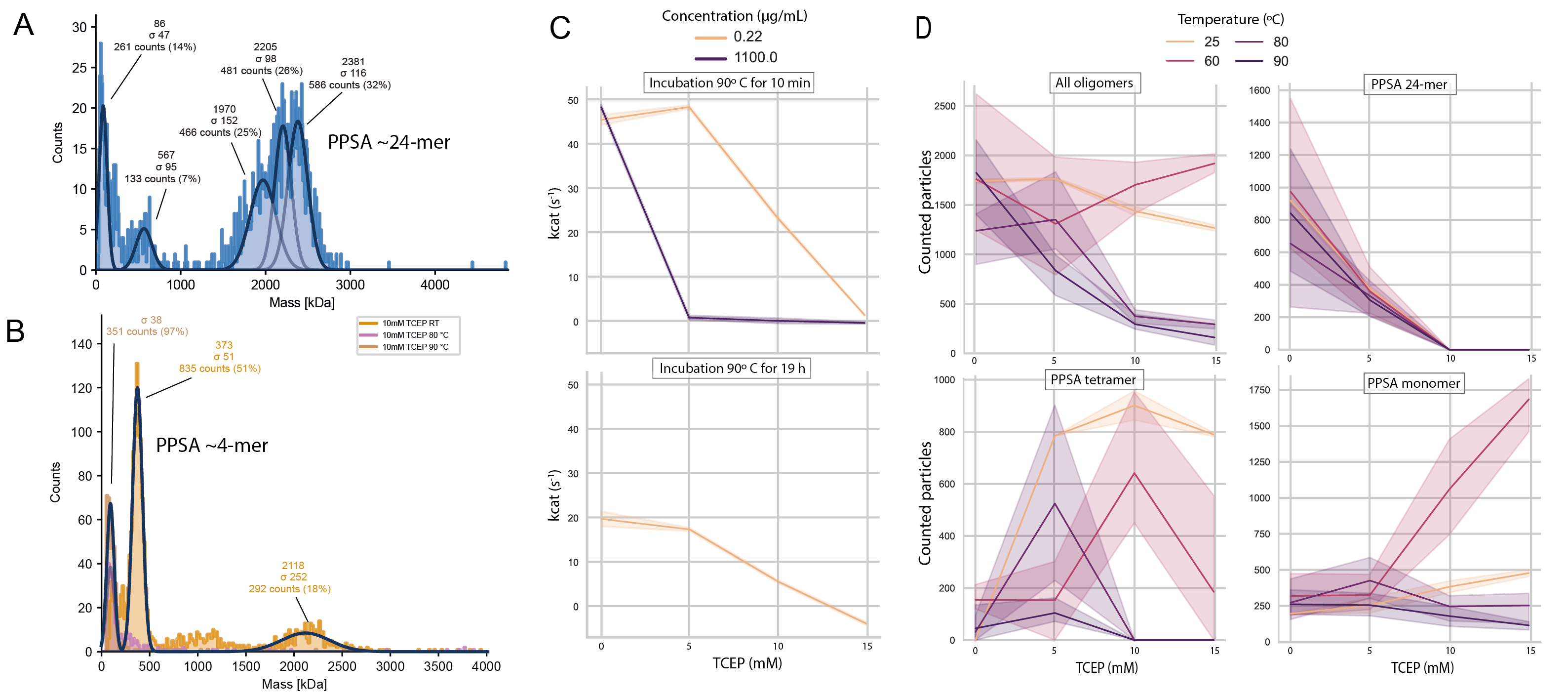

**Supplementary Figure 3. The PPSA tetrameric sub-complex is the functional unit but is not thermo stable.** The purified PPSA fraction, although highly pure, is an ensemble of >2 MDa complexes as measured by mass photometry **(A)**. Functional assays were conducted in duplicate after 10 min incubation at the given concentration **(B)**. Mass photometry measurements at the different TCEP concentrations were conducted from the same starting sample from three independent replicates and incubated at 10 µg/mL at the different temperatures/TCEP concentrations **(C,D)**. The number of particles within a given mass range were assigned to a specific oligomerization state of PPSA (see Methods for details)

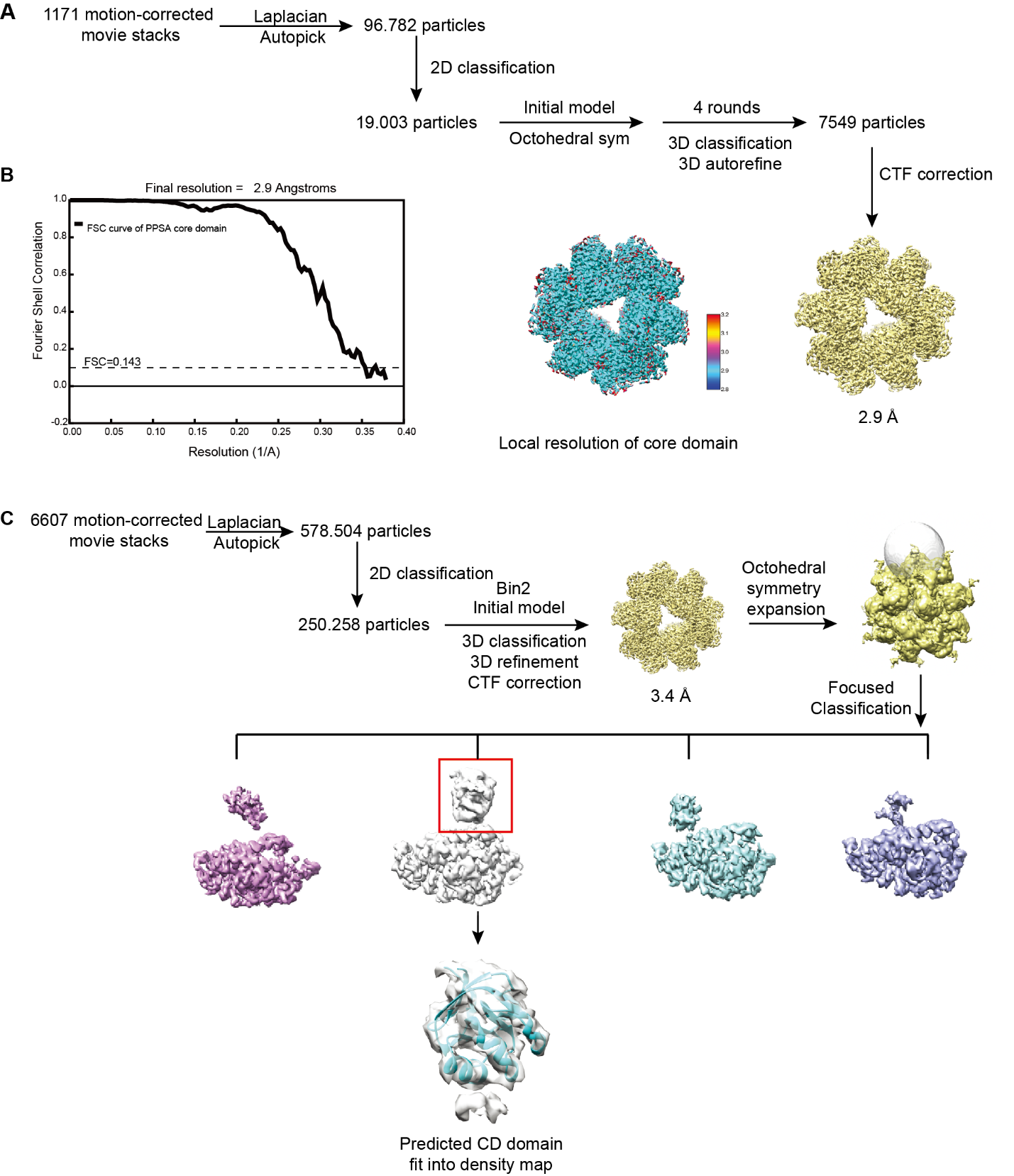

**Supplementary Figure 4. Cryo-EM SPA of the PPSA core.** Cryo-EM data processing workflow of the PPSA PBD domain and its local resolution **(A)**. FSC curve of the PPSA PBD domain **(B)**. Cryo-EM data processing of the PPSA CD domain **(C)**.

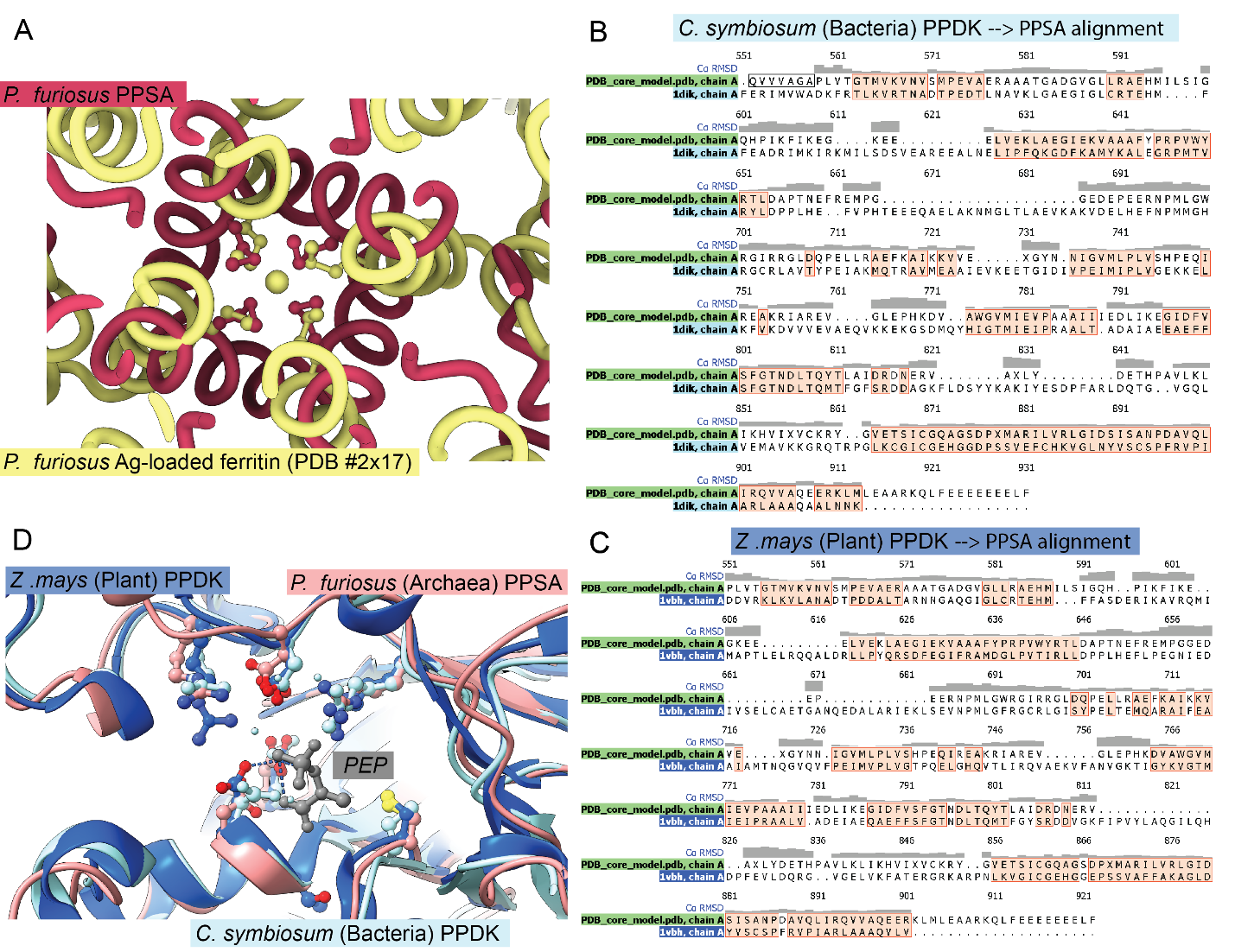

**Supplementary Figure 5**: **Conserved structural features of the PBD core**. Structural similarities between the Met-Fe(II) cluster of PPSA PBD core tetramers and the Met-Ag cluster of ferritin (PDB #2x17), both from *P. furiosus* **(A)**. Structural alignment of Bacterial (PDB #1dik) **(B)** and Eukaryotic (PDB #1vbh) **(C)** PPDK and the PBD core model from *P. furiosus*. The key residues facing the catalytic PBD core, conserved in the 3 kingdoms of life, are shown along with PEP positioning from PDB #1vbh **(D)**.

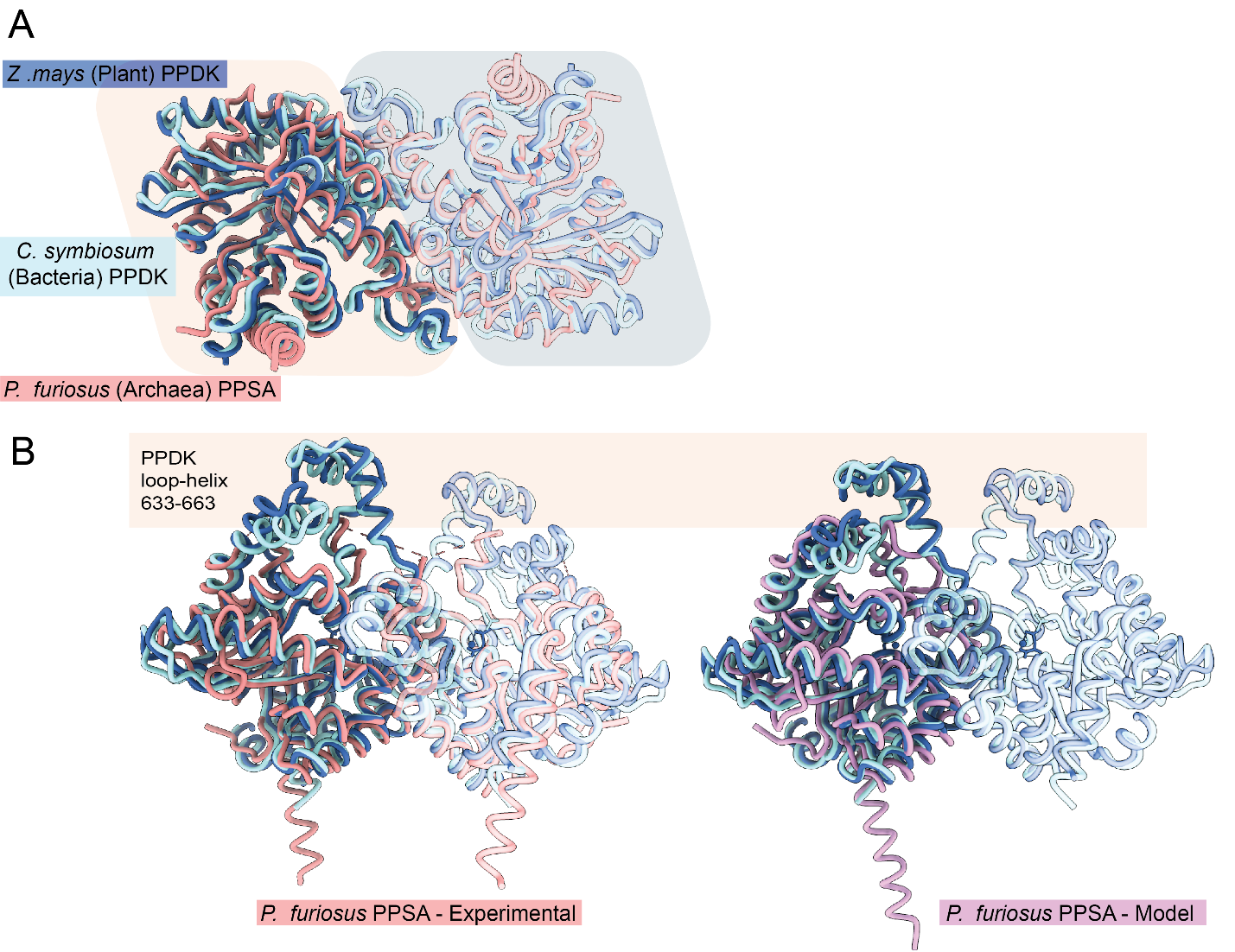

**Supplementary Figure 6. Structural conservation of the dimeric interface of the PBD in mesophilic PPDK enzymes of Eukaryotes and Prokaryotes.** Structural alignment of Bacterial (PDB #1dik) and Eukaryotic (PDB #1vbh) PPDK and the PBD core model from *P. furiosus* in top-view **(A)** and side-view **(B)**. Both the cryo-EM derived model (left) and the predicted model (right) are superimposed to the Bacterial (PDB #1dik) and Eukaryotic (PDB #1vbh) PPDK, with the additional 2 helix motif in the PPDK enzymes highlighted.

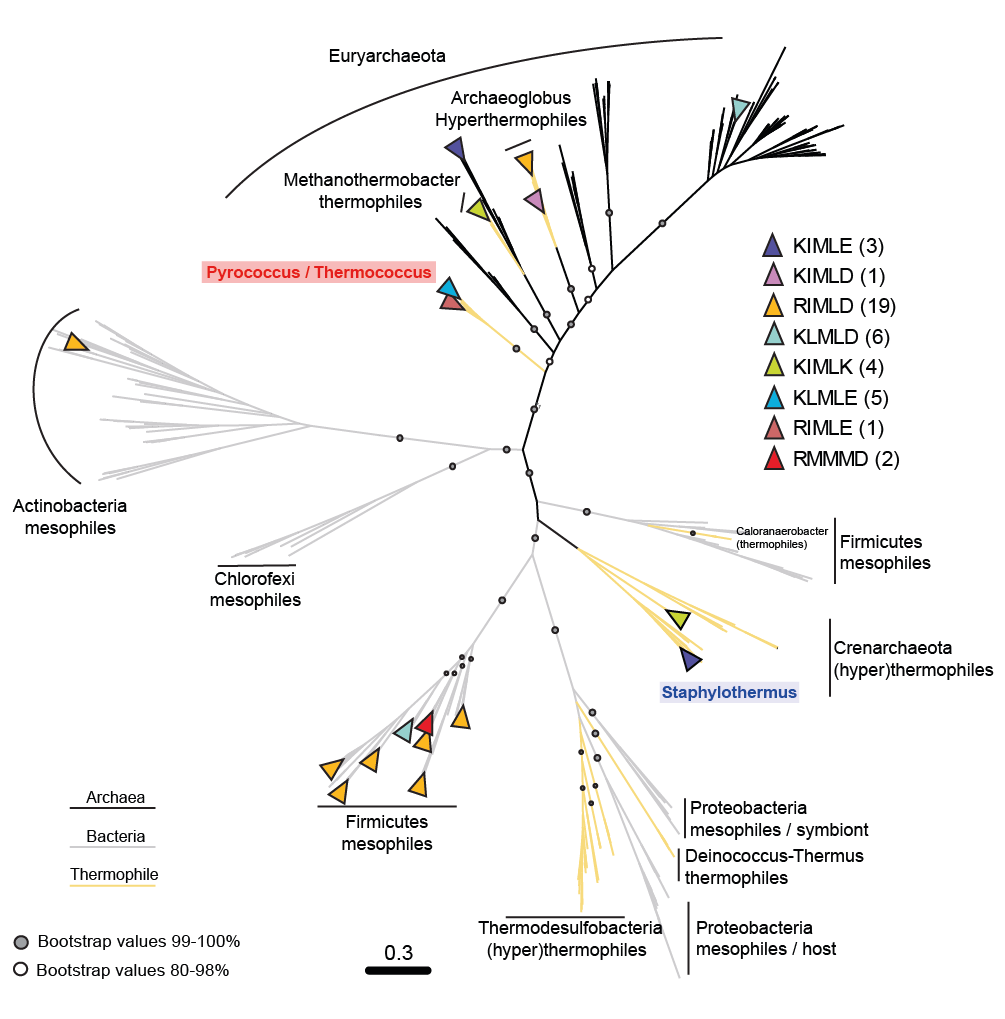

**Supplementary Figure 7. Phylogenetic analysis of PPSA homologs suggests an ancient origin of the 24-meric state common to hyperthermophiles.** Unrooted maximum likelihood phylogeny (RAxML, LG+C60) based on 325 homologous PPSA sequences from Archaea and Bacteria species. Node supports are indicated as filled or empty circles if bootstrap support (200 replicates) is >99% or 80-98%, respectively. The scale bar represents the average number of substitutions per site. Archaea (black lines) and Bacteria (grey lines) branches are distinguished, with known thermophilic species from both kingdoms highlighted in yellow. Motifs indicating a probable 24-meric assembly are indicated as color-coded triangles. The two taxa where this oligomerization is known (*Pyrococcus* in red and *Staphylothermus*(Li et al., 2000) in blue) are highlighted.

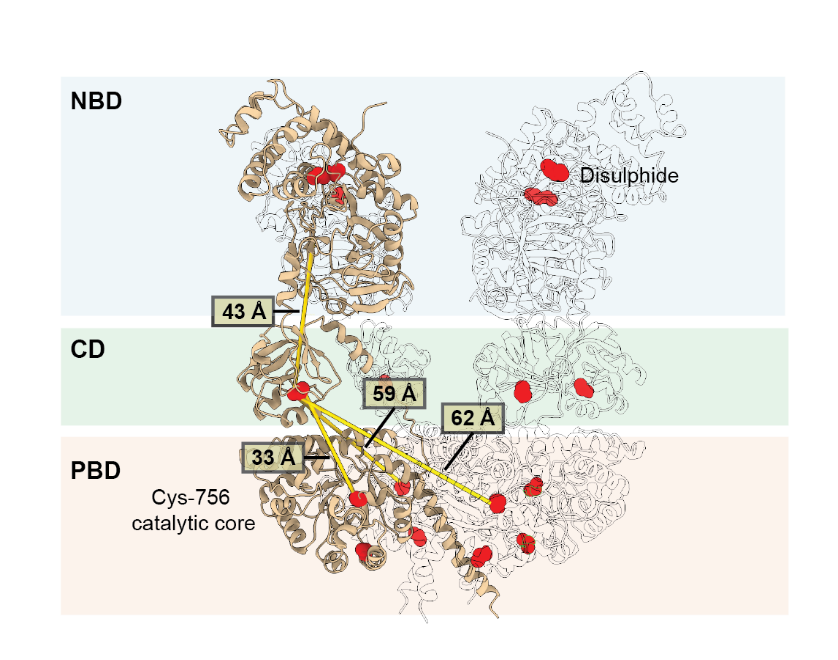

**Supplementary Figure 8. Distance between catalytic/functional domains of PPSA and cysteine positions.** The approximate distance between catalytic domains of the (c1)_4_ conformation is shown as the minimal distance with nearest neighboring subunits (or within the same subunit). The distance between CD and NBD domains is based on the modelled structure, while CD-PBD distance is based on the experimental model and the catalytic Cys-756 position. The positions of the 5 cysteines are highlighted in red, showing the spacing hindering the possible formation of inter-chain disulfide bonds.

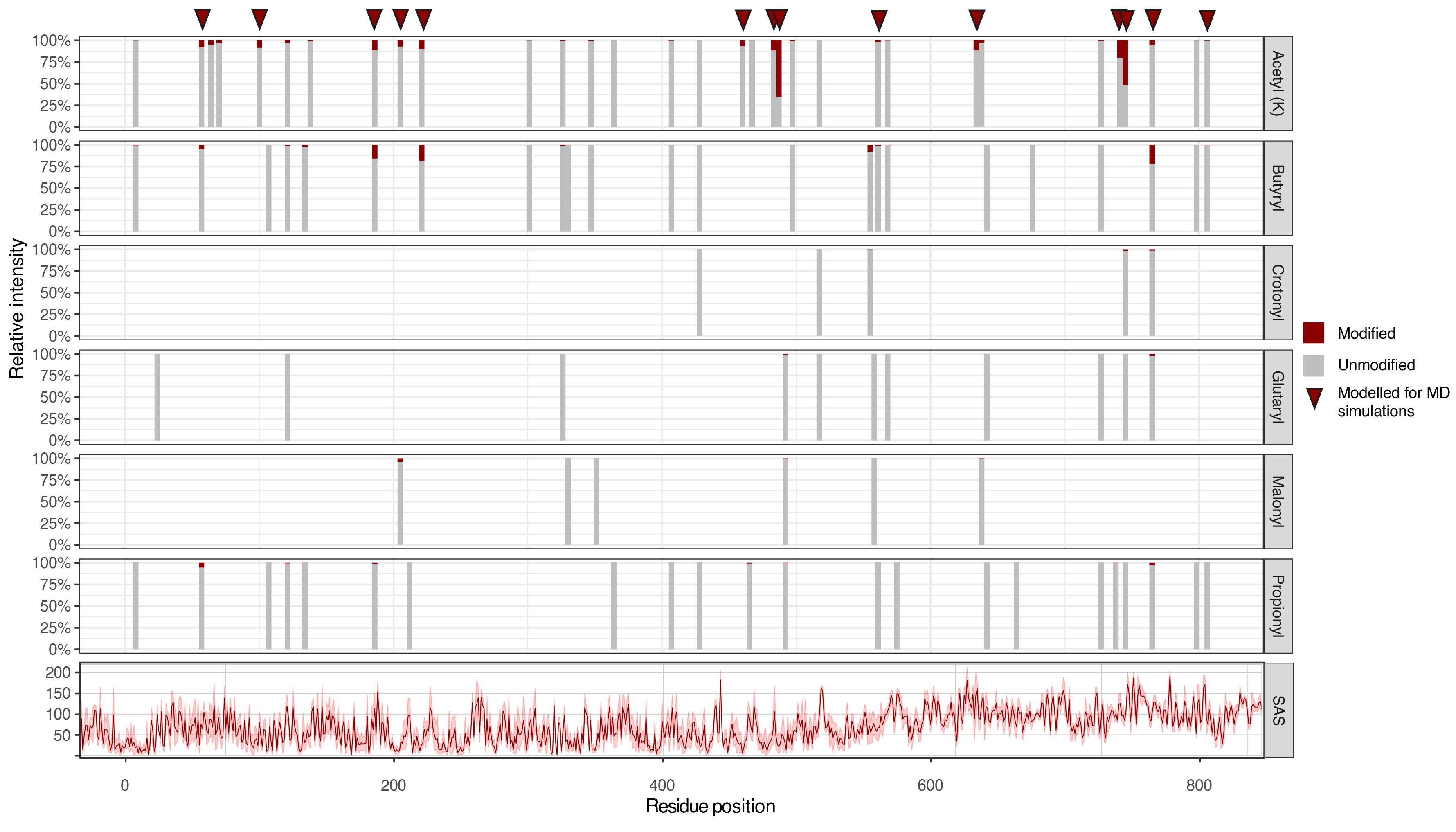

**Supplementary Figure 9. Location and relative abundance of various lysine acylation modification sites on PPSA.** Acetylation stoichiometry is depicted as a normalized intensity ratio between modified (red) and unmodified (grey) peptides. Only one modification is allowed per peptide (per modification), therefore the depicted stoichiometry is an underestimation given the heterogeneity of site-specific modifications. The Solvent Accessible Surface (SAS) in Å was calculated on 8 PPSA monomers after extensive MD simulations (see the appropriate Results section) using the PISA web service (<https://www.ebi.ac.uk/msd-srv/prot_int/cgi-bin/piserver>). The acK sites selected for MD simulations are marked with a triangle.

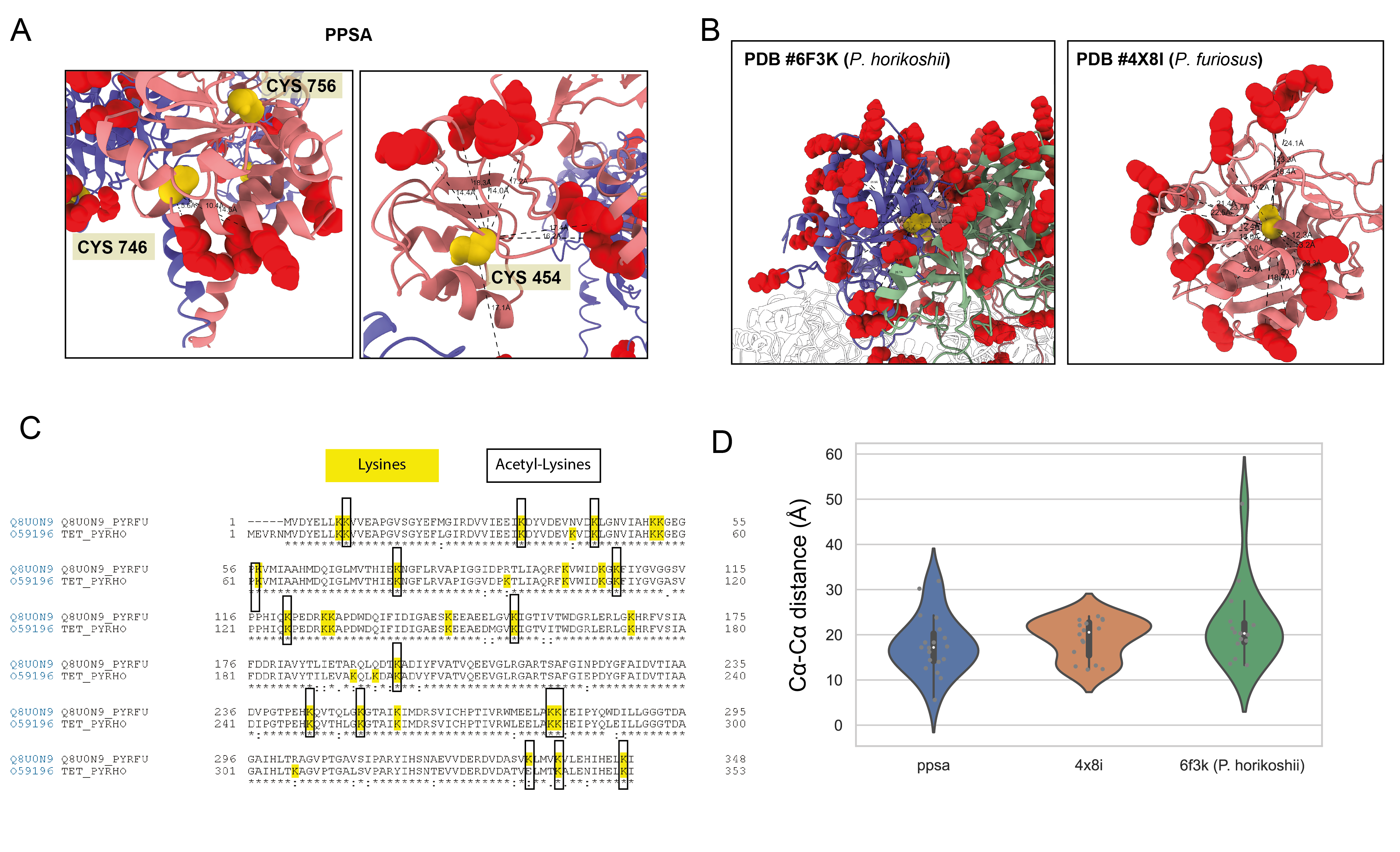

**Supplementary Figure 10: Cysteine and acetylated lysine residues proximity as a mechanism for non-enzymatic lysine acetylation.** The distances between “free” cysteine and acetylated lysine residues are displayed and measured for the PBD and CD domains of PPSA **(A)** of the lysyl aminopeptidase from *P. furiosus* (PDB #4x8i) and the aminopeptidase TET2 from *P. horikoshii* (PDB #6f3k) **(B)**. The latter showing 89% sequence identity with the *P. furiosus* homolog (Q8U0N), and for 15 of the 16 detected acK residues **(C)**. The average distance between the cysteine and acK residues in these complexes is consistently ~20 Å **(D).**

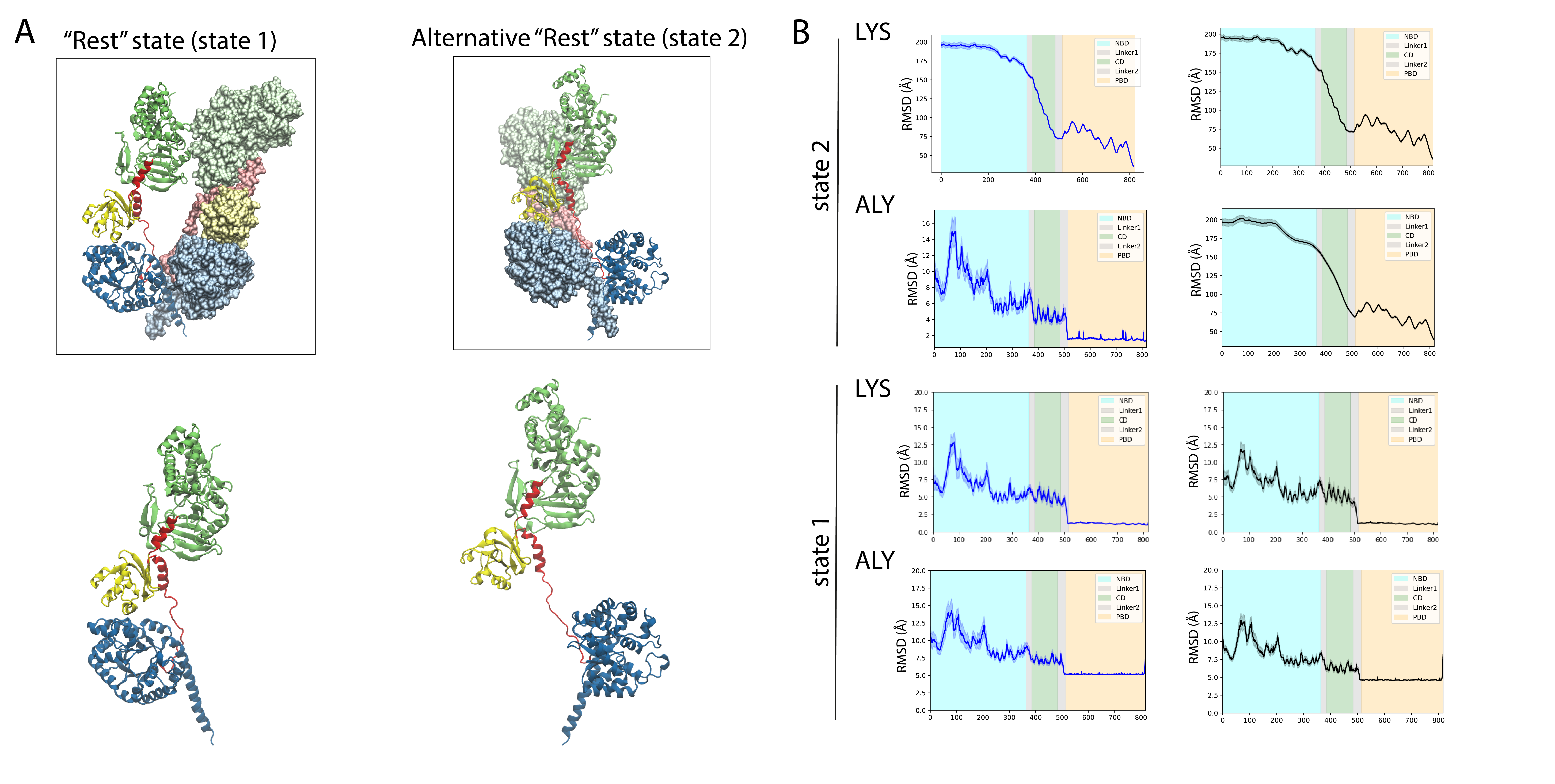

**Supplementary Figure 11. Two alternative conformations of PPSA tetramers in the “rest” state were simulated over 125ns.** The two alternative configurations possible (state 1 and state 2) considering the long CD-PBD connecting loop were modeled **(A)** and tested for stability over a 125 ns MD simulation runs in duplicate **(B)**. The root mean square deviation (RMSD) from the initial structure highlights how state 2 is instable, independently from the presence or absence of acetylation.

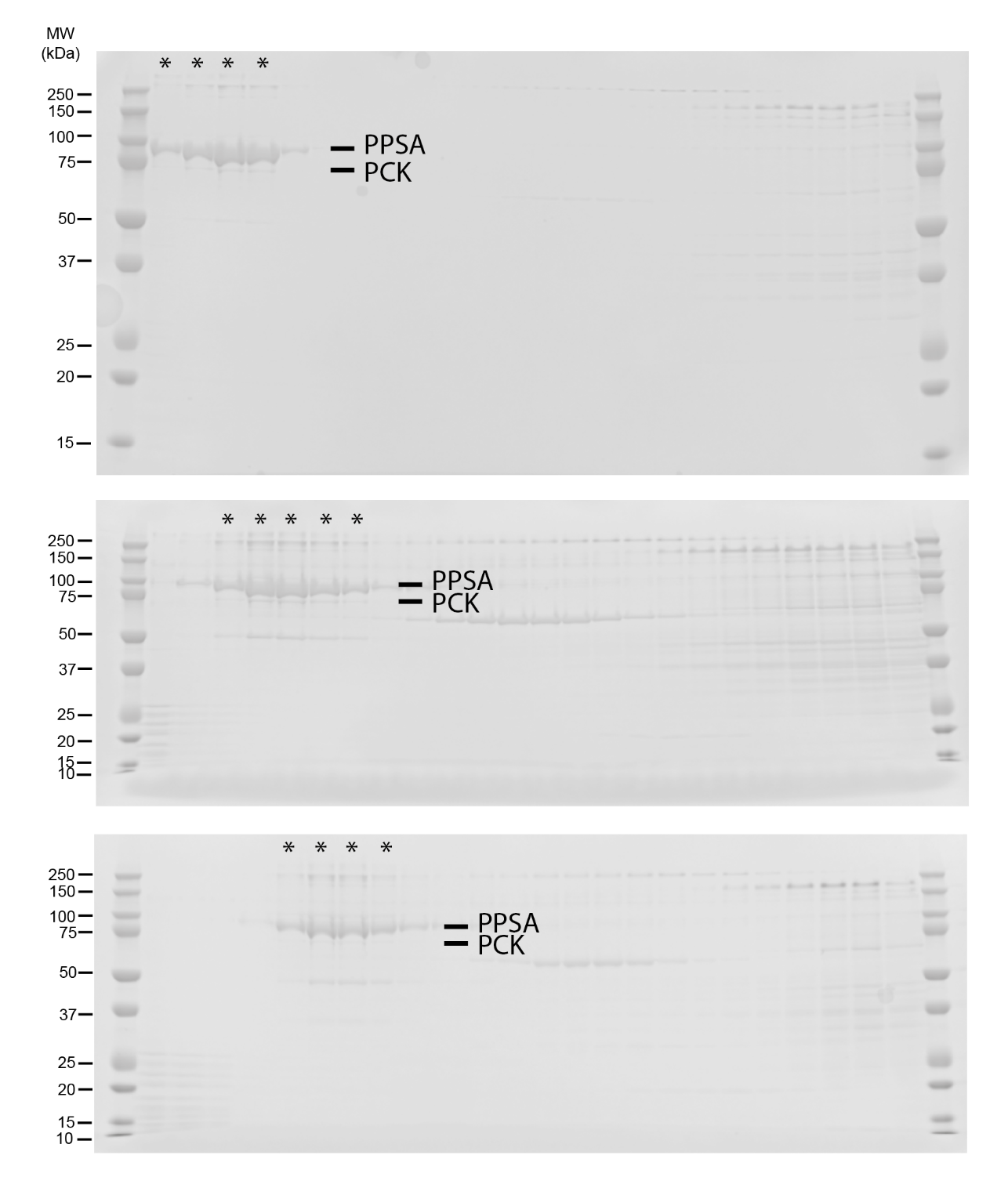

**Supplementary Figure 12. PCK is stably interacting with PPSA 24-mer.** SDS-PAGE profile of sucrose gradient fractions from three independent rounds of PPSA purification showing the reproducible co-migration of the ~70 kDa PCK interactor. Bands relatively quantified with densitometry measurements shown in Figure 5A are indicated with an asterisk (*).

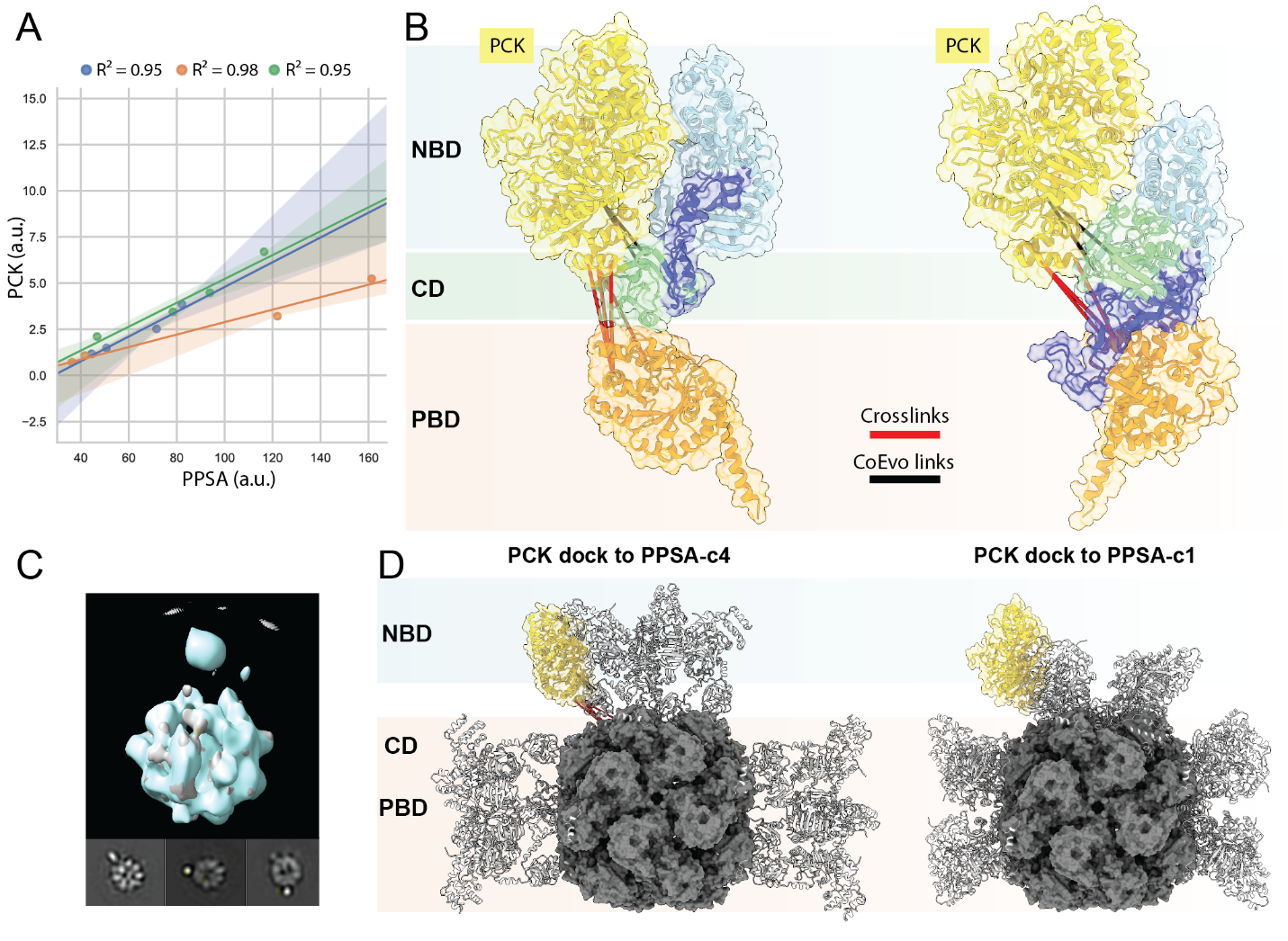

**Supplementary Figure 13. Stable association of an adjuvant enzyme challenges the homomeric nature of PPSA complex.** Consistent comigration of Phosphoenolpyruvate carboxykinase (PCK, ~70KDa) and PPSA was evaluated by SDS-PAGE (Supplementary Figure 10) and densitometry measurements from three independent purification rounds. Density of the PCK band against the PPSA band of 4 consecutive 150 µL fractions from the sucrose gradient are plotted, linear regression curves interpolating 4 data point are shown **(A)**. Molecular docking of Phosphoenolpyruvate carboxykinase (PCK) to the c4 and c1 conformations on the full 24-mer (other subunits are not displayed for clarity) using distance restraints from XL-MS and co-evolution analysis **(B)**. Direct comparison with low resolution cryo-EM maps and relative projections of 2D class averages **(C)** and overall positioning of the docked PCK within a fully assembled 24-meric PPSA (the cryo-EM derived core is represented with its molecular surface) **(D)**.

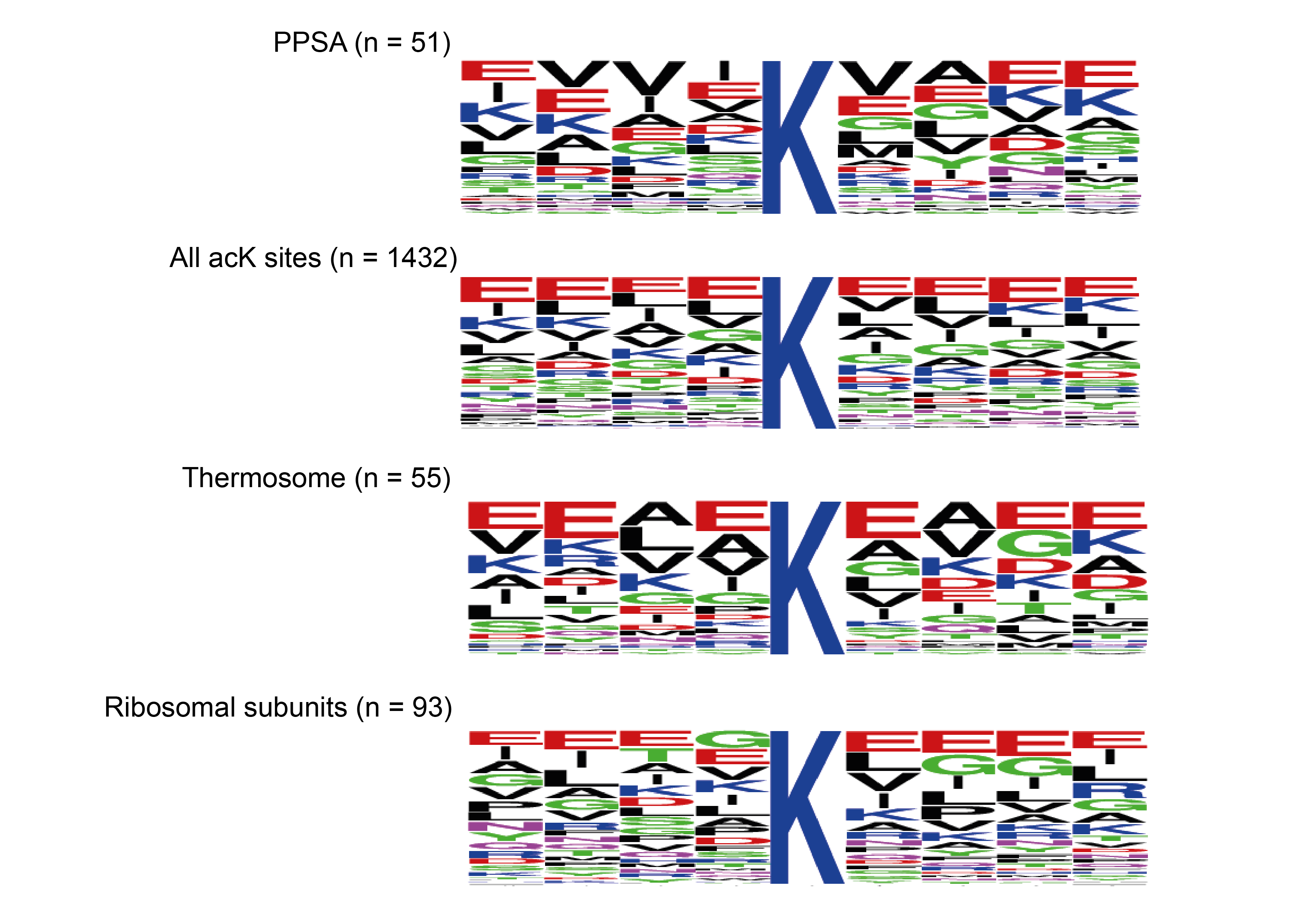

**Supplementary Figure 14. Sequence motif analysis of regions flanking the acK sites.** Sequence motif analysis for acetylated lysine residues using the flanking 6 amino acids (+6 and -6) of the 51 acetylated lysines of PPSA, and compared with all detected acK sites. The same analysis for one homomeric comples (Thermosome) and one heteromeric complex (Ribosome) are shown.

**Supplementary Note 1**

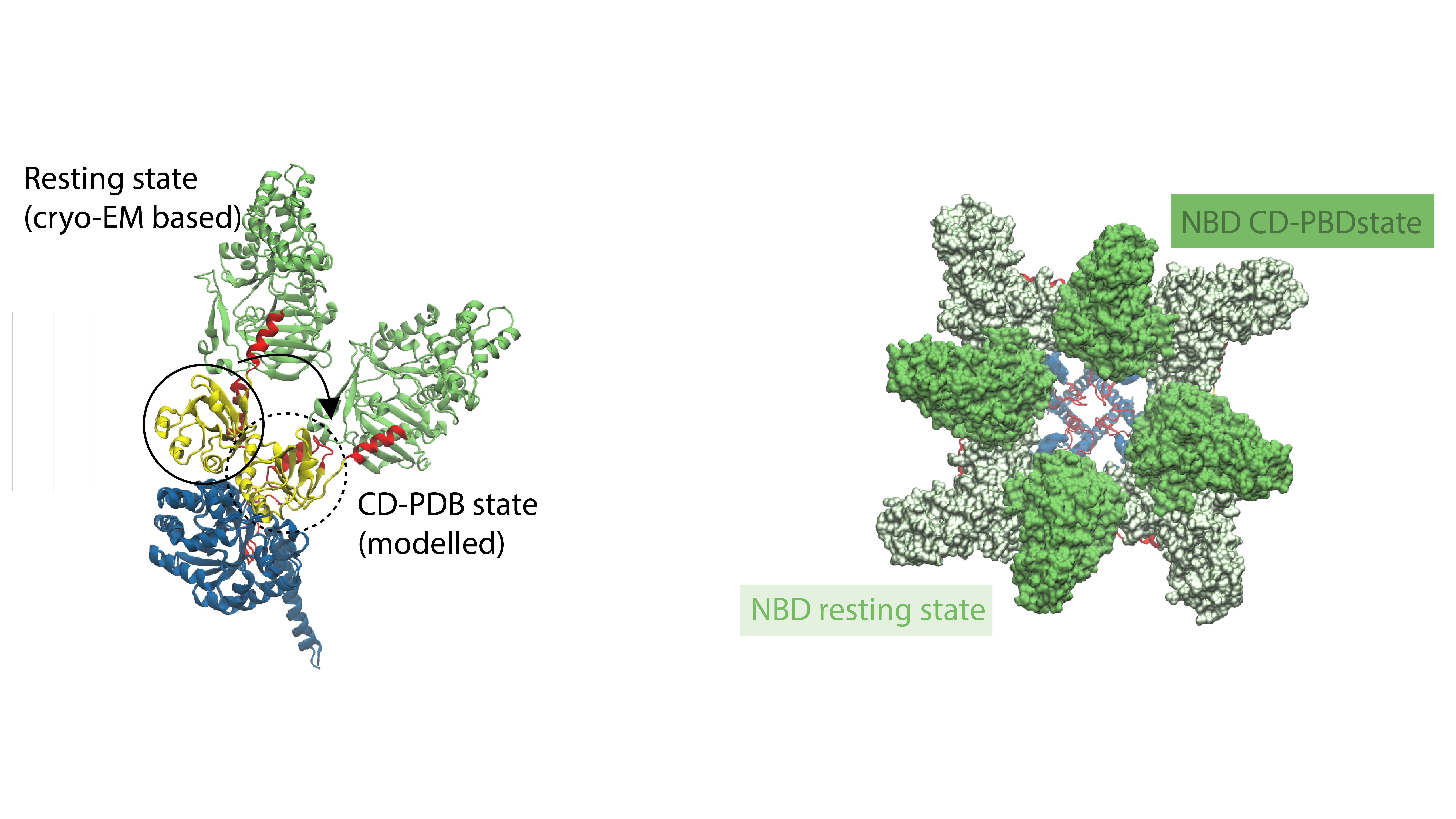

The four predicted PPSA states suggest a catalytic mechanism that takes place in these PPSA tetramers. Besides including the performance of two reaction mechanisms similar to PPDK, the models also indicate an interplay between neighboring PPSA monomers within each tetramer in which two PPSA proteins are required to perform the complete catalytic cycle (PPSA1 and PPSA2), a unique mechanism compared to formerly resolved PPDK structures in which one PPDK protein performs the complete cycle (Lim et al., 2007). The modelled c1 state represents an “inactive” state of PPSA1 in the PPSA tetramer at the beginning of the ATP/pyruvate conversion cycle in which CD is interacting with NBD and does not form residue-residue contacts with PBD. This potentially is a conformation in which NBD can start interacting with ATP to initiate ATP-to-AMP conversion, releasing a phosphate moiety that covalently binds to CD. In the c2 conformation, phosphorylation of CD (PPSA1) is assumed and CD has undergone a large conformational change, repositioning CD (PPSA1) at the active site of PBD of PPSA2 in such a way that pyruvate can be bound to the PBD site. Hence, the phosphorylated region on CD (PPSA1) is in close proximity with the conserved active site on PBD (PPSA2), generating a conformation for a potential pyruvate-PEP conversion(Lim et al., 2007). The c3 state is similar to c2 as CD of PPSA1 is interacting with PDB of PPSA2 as well. However, in c3 the interaction between CD and NBD is enhanced by the formation of a helical region in the NBD-CD linker while the interaction between CD (PPSA1) and PDB (PPSA2) remains unaltered. This helix formation in the NBD-CD linker prepares the PPSA structure for the c4 state in which the complete CD-NBD region is relocated on top of PPSA’s PBD. This c4 conformation resembles the PPSA conformation found in the cryo-EM data the most, in which CD (PPSA1) is located on top of PBD (PPSA2) without including an interaction between the phosphorylated region on CD and the active site on PBD. The c1, c2 and c3 states can be combined within a tetramer whereas the c4 conformation requires all PPSA units to be in this conformation, alluding to a role for allostery within this catalytic cycle. The structural model most in line with the cryo-EM data is the “(c4)_4_” arrangement in which the CD domain is not functionally associated to NBD or PBD, this conformation will therefore be referred to here as the “rest” position. The linkers between NDB-CD and CD-PBD appear to play a crucial role in relocating CD and NBD, depending on the step in the catalytic cycle. Even though the presence of these flexible regions is not surprising for mesophilic organisms, it is likely a liability at the extreme conditions (100 ˚C) in which this enzyme operates due to a potential increase in dynamics induced by the elevated temperatures.
